# Plastid phylogenomics reveals evolutionary relationships in the mycoheterotrophic orchid genus *Dipodium* and provides insights into plastid gene degeneration

**DOI:** 10.1101/2024.02.05.578113

**Authors:** Stephanie Goedderz, Mark A. Clements, Stephen J. Bent, James A. Nicholls, Vidushi S. Patel, Darren M. Crayn, Philipp M. Schlüter, Katharina Nargar

## Abstract

The orchid genus *Dipodium* R.Br. (Epidendroideae) comprises leafy autotrophic and leafless mycoheterotrophic species, the latter confined to sect. *Dipodium*. This study examined plastome degeneration in *Dipodium* in a phylogenomic and temporal context. Whole plastomes were reconstructed and annotated for 24 *Dipodium* samples representing 14 species and two putatively new species, encompassing over 80% of species diversity in sect. *Dipodium*. Phylogenomic analysis based on 68 plastid loci including a broad outgroup sampling across Orchidaceae found sect. *Leopardanthus* as sister lineage to sect. *Dipodium. Dipodium ensifolium*, the only leafy autotrophic species in sect. *Dipodium* was found sister to all leafless, mycoheterotrophic species, supporting a single evolutionary origin of mycoheterotrophy in the genus. Divergence time estimations found that *Dipodium* arose ca. 33.3 Ma near the lower boundary of the Oligocene and crown diversification commenced in the late Miocene, ca. 11.3 Ma. Mycoheterotrophy in the genus was estimated to have evolved in the late Miocene, ca. 7.3 Ma, in sect. *Dipodium*. The comparative assessment of plastome structure and gene degradation in *Dipodium* revealed that plastid *ndh* genes were pseudogenised or physically lost in all *Dipodium* species, including in leafy autotrophic species of both *Dipodium* sections. Levels of plastid *ndh* gene degradation were found to vary among species as well as within species, providing evidence of relaxed selection for retention of the NADH dehydrogenase complex within the genus. *Dipodium* exhibits an early stage of plastid genome degradation as all species were found to have retained a full set of functional photosynthesis-related genes and housekeeping genes. This study provides important insights into plastid genome degradation along the transition from autotrophy to mycoheterotrophy in a phylogenomic and temporal context.

## 1 Introduction

Heterotrophic plants - plants that rely on other organisms for energy and nutrients - are remarkable survivors, exhibiting often curious morphological, physical, or genomic modifications, reflecting evolutionary relaxed selective pressure on photosynthetic function (Graham et al., 2017; Barrett et al., 2019). Advances in next generation sequencing and bioinformatic pipelines have vastly accelerated the characterisation of plastid genomes (plastomes), including of heterotrophic plants, providing new insights into plastome evolution. Plastomes of heterotrophic plants often exhibit greatly altered structure and gene content due to photosynthesis-related genes that are no longer required (Delannoy et al., 2011; Barrett et al., 2014; Lam et al., 2015; Graham et al., 2017; Braukmann et al., 2017; Barrett et al., 2018; Wicke and Neumann 2018; Qu et al., 2019; Barrett et al., 2019; Klimpert et al., 2022; Peng et al., 2022; Wen et al., 2022). Hence, heterotrophic plants offer excellent opportunities to gain insight into plastome evolution under relaxed selection.

Early non-phylogenomic studies on plastome evolution in heterotrophic plants allowed the discovery of large-scale plastome evolutionary patterns and, moreover, stimulated research into fine-scale, phylogenetic comparative approaches (e.g., Delannoy et al., 2011; Logacheva et al., 2011; Roma et al., 2018). Thus far, most phylogenetic comparative studies included plastomes of taxa scattered across families, tribes, or genera (e.g., Kim et al., 2015; Feng et al., 2016; Niu et al., 2017; Lallemand et al., 2019; Li et al., 2020; Kim et al., 2020; Tu et al., 2021; Kim et al., 2023). Yet, phylogenetic, comparative approaches at infrageneric level are still scarce (e.g., Barrett et al., 2018; Barrett et al., 2019).

Orchidaceae, one of the two largest flowering plant families, has undergone a greater number of independent transitions from autotrophy to heterotrophy than any other land plant lineage (Merckx 2013; Christenhusz and Byng 2016; Jacquemyn and Merckx 2019). The family comprises several heterotrophic orchid lineages which rely to some extent on mycorrhizal fungi for carbon and other nutrients i.e., initial, partial, or full mycoheterotrophy (Merckx 2013).

So far, most examined mycoheterotrophic orchid plastomes exhibited degradation patterns similar to those found in heterotrophic plastomes of other plants. These include a reduction in genome size, decrease in guanine-cytosine (GC) content, rearrangements, pseudogenisations and gene losses (e.g., Delannoy et al., 2011; Barrett et al., 2018; Lallemand et al., 2019; Barrett et al., 2019; Wen et al., 2022). Moreover, whole plastome sequencing has revealed patterns of plastid gene degradation for various heterotrophic plastomes which led to the development of conceptual models to predict the evolutionary transition from autotrophy to heterotrophy of the plastid organelle (e.g., Graham et al., 2017; Barrett et al., 2019). Several studies in mycoheterotrophic orchid lineages found support for these models which predict a progression from losses of the chloroplast *ndh* genes to genes encoding complexes which are directly involved in photosynthesis (e.g., *psa*, *psb*) to more general ‘housekeeping’ genes (e.g., *acc*D, *mat*K) (Wicke and Naumann 2018; Barrett et al., 2018; Barrett et al., 2019; Kim et al., 2020; Kim et al., 2023).

Interestingly, degraded *ndh* genes were also found in some autotrophic orchids (e.g., Kim et al., 2015; Niu et al., 2017; Kim and Chase 2017; Lallemand et al., 2019; Kim et al., 2023). This appears curious, as the *ndh* genes encode proteins of the NADH dehydrogenase complex (NDH complex) which is assumed to play a role in cyclic electron flow and thus fine-tunes photosynthesis (Yamori et al., 2015; Peltier et al., 2016). Degradation of *ndh* genes is hypothesised to have led to additional structural changes of the plastome (Kim et al. 2015). In particular, *ndh*F gene loss was correlated with shifts in the position of the junction of the inverted repeat/small single copy (IR/SSC) region in Orchidaceae and other plants (Kim et al., 2015; Niu et al., 2017; Dong et al., 2018; Roma et al., 2018; Thode and Lohmann 2019; Li et al., 2021; Könyves et al., 2021). However, within Orchidaceae, degradation of *ndh* genes was found to vary even among closely related species (e.g., Kim et al., 2015; Feng et al., 2016; Kim and Chase, 2017; Barrett et al., 2018; Barrett et al., 2019) which suggests the genes for the NDH complex may be under relaxed selective pressure in several orchid lineages (Kim and Chase, 2017). Moreover, previous studies found that *ndh* degradation patterns vary considerably and have been independently degraded among orchids (Kim et al., 2015; Niu et al., 2017; Kim and Chase 2017; Lallemand et al., 2019).

The orchid genus *Dipodium* R.Br. (Cymbidieae) contains both autotrophic and mycoheterotrophic species and thus represents a suitable model system in which to address hypotheses of plastome evolution. The genus comprises 39 species and is divided into two sections, *Dipodium* and *Leopardanthus* (Blume) O. Kuntze, based on morphological and geographical evidence (O’Byrne, 2017; Jones, 2021). Sect. *Leopardanthus* (26 species) is distributed in the floristic regions of Malesia and Australasia (O’Byrne, 2017). All species of sect. *Leopardanthus* are green leafy plants and non-uniform in habit (O’Byrne, 2017). Section *Dipodium* occurs predominantly in Australasia, with nearly all species being endemic in Australia. One species occurs in New Guinea (*D. elatum* J.J.Sm.), one species extends into the Pacific region (*D. squamatum* (G.Forst.) Sm. (New Caledonia and Vanuatu), and one occurs in Malesia (*D. gracile* Schltr. (Sulawesi) (Schlechter, 1911; O’Byrne, 2017; POWO, 2023; WFO, 2023). In contrast to sect. *Leopardanthus*, most species of sect. *Dipodium* are non-climbing terrestrials, forming subterranean rhizomes and erect flowering stems with highly reduced, non-photosynthetic leaves (i.e., scales) (**Figure 1**, B). Hence, species within sect. *Dipodium* are generally assumed to be fully mycoheterotrophic (O’Byrne, 2014). However, one Australian species of sect. *Dipodium*, *D. ensifolium* F.Muell., stands out as a leafy terrestrial (**Figure 1**, A).

**Figure 1:**
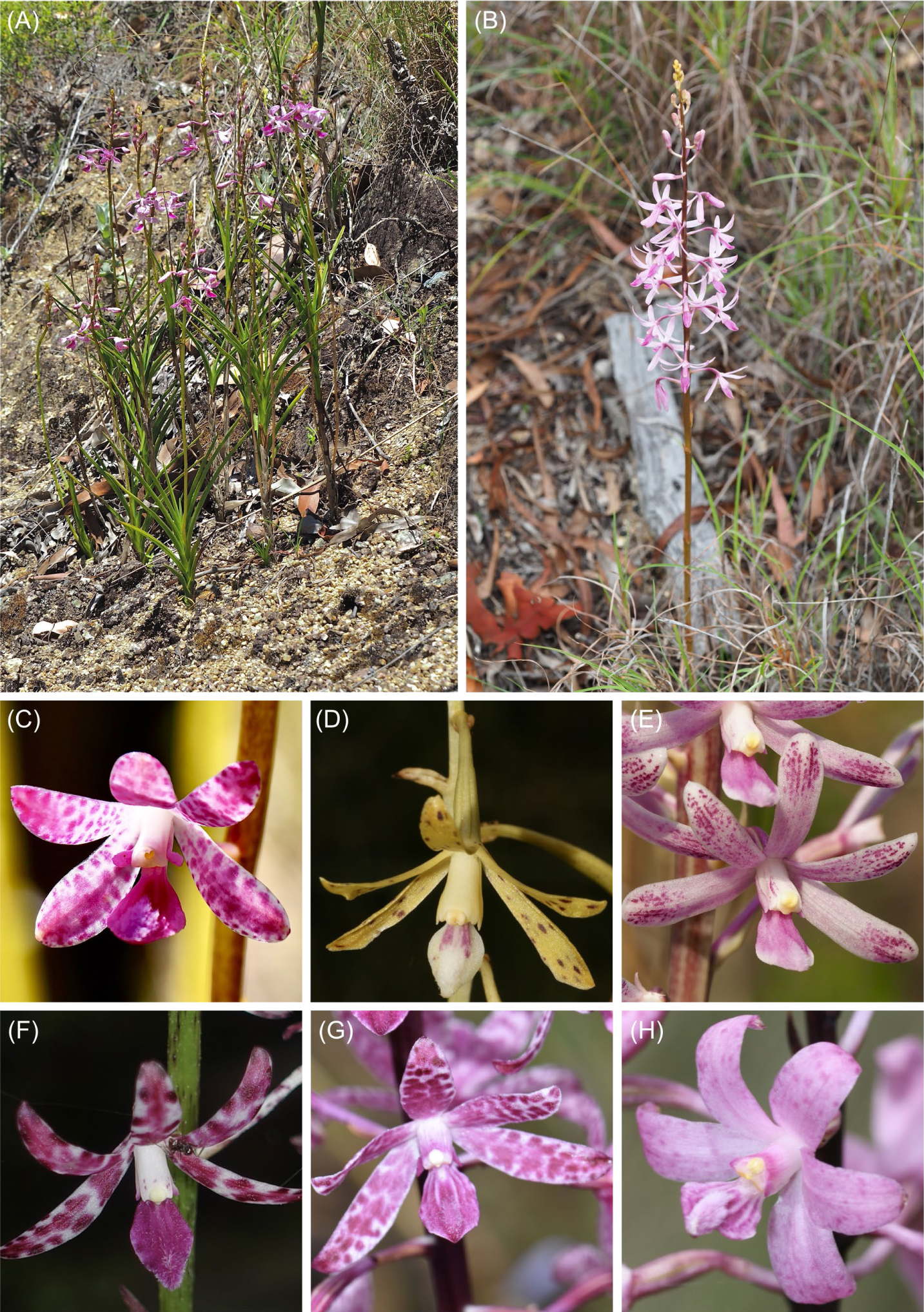
Habit and flowers of *Dipodium* sect. *Dipodium*. A. *D. ensifolium;* B. *D. elegantulum* (note the green to purplish inflorescence stem); C. *D. ensifolium*; D. *D. interaneum*; E. *D. elegantulum;* F. *D. variegatum*; G. *D. punctatum*; H. *D. roseum*. (Photos: A-C, E: S. Goedderz; D, F -H: M.A. Clements.)

The aims of this study were to:

1. sequence and assemble plastid genomes for species of *Dipodium* to elucidate patterns of plastid genome modification (e.g., rearrangement, structural variation, pseudogenisation, gene loss) across autotrophs and mycoheterotrophs within the genus and examine gene degradation in context of current models of plastome degradation in heterotrophic plants.
2. infer phylogenomic relationships within section *Dipodium* and among closely related autotrophic relatives (i.e., *Dipodium* section *Leopardanthus*).
3. estimate divergence times of *Dipodium* to assess the origin of mycoheterotrophy within the genus and elucidate over which evolutionary timeframes plastid gene degradation and losses has taken place within *Dipodium*.

## 2 Material and methods

### 2.1 Plant material

For this study, we sampled all known Australian species of section *Dipodium* and one representative of section *Leopardanthus* (**Table 1**). Based on previous molecular systematic studies (Serna-Sánchez et al., 2021; Pérez-Escobar et al. 2023; Zhang et al. 2023), an extended outgroup from closely related orchid genera within subtribe Eulophiinae (*Eulophia* R.Br., *Geodorum* Andrews) and subtribe Cymbidiinae (*Cymbidium* Sw., *Acriopsis* Reinw. ex. Blume) was sampled (**Table 1**). Specimens studied were from different regions within Australia, with the exception of one specimen from Papua New Guinea (*D. pandanum 2*) (**Table 1**).

**Table 1.**
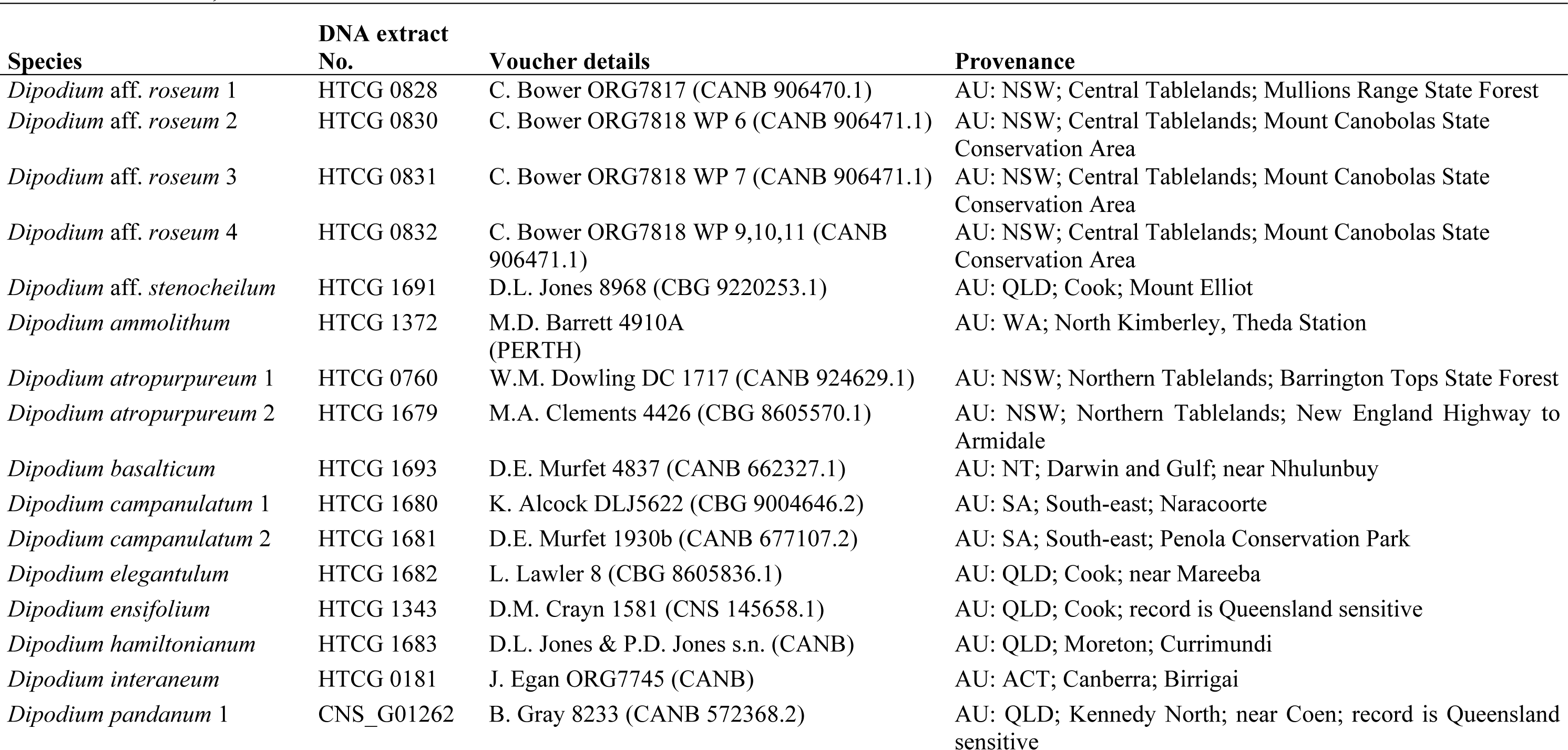
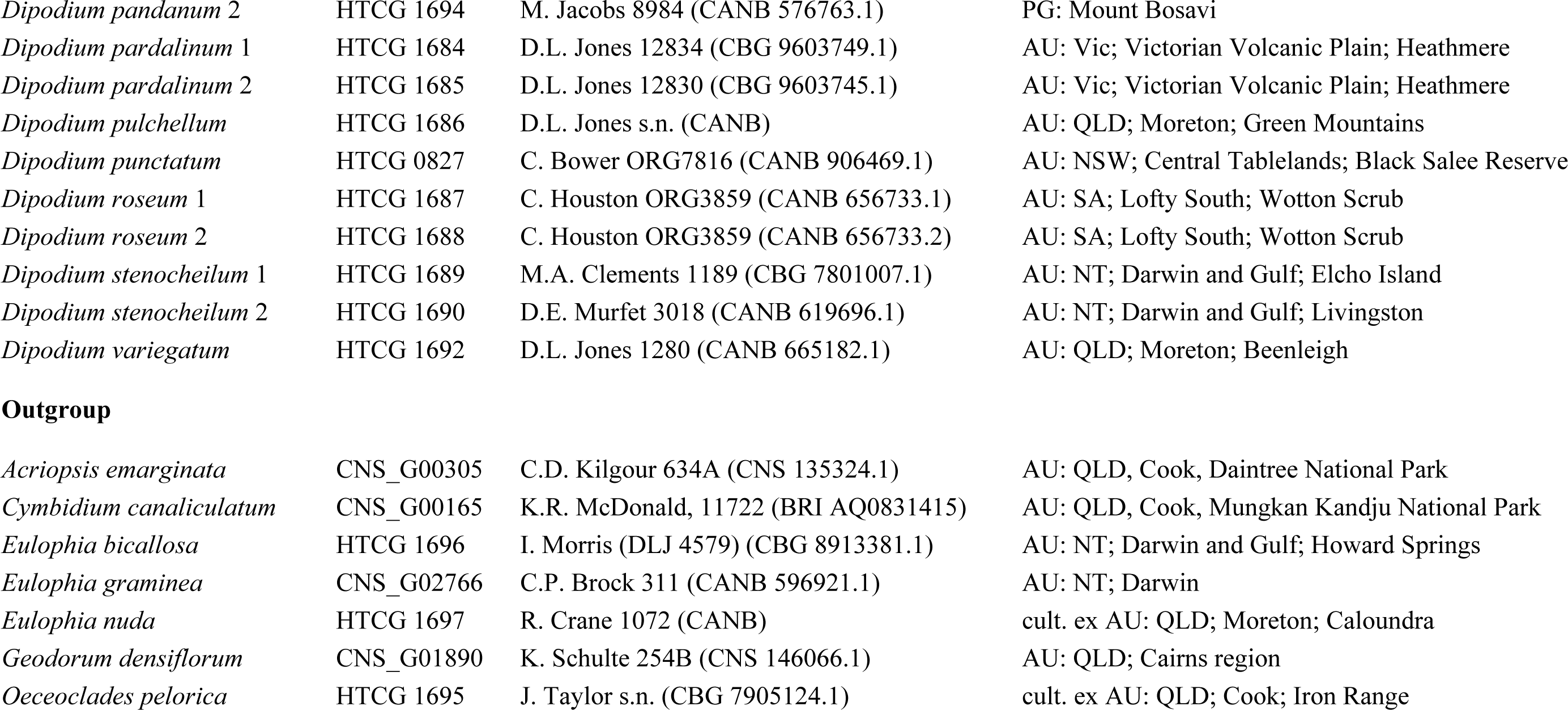
Plant material used in this study inclusive voucher details and provenances with botanical districts. Taxonomy according to the Australian Plant Census (APC, 2023). CANB = Australian National Herbarium, CNS = Australian Tropical Herbarium. AU = Australia, PG= Papua New Guinea. ACT = Australian Capital Territory, NT = Northern Territory, NSW = New South Wales, SA = South Australia, QLD = Queensland, WA <u>= Western Australia, Vic = Victoria.</u>

| Species | DNA extract No. | Voucher details | Provenance |
| --- | --- | --- | --- |
| *Dipodium* aff. *roseum* 1 | HTCG 0828 | C. Bower ORG7817 (CANB 906470.1) | AU: NSW; Central Tablelands; Mullions Range State Forest |
| *Dipodium* aff. *roseum* 2 | HTCG 0830 | C. Bower ORG7818 WP 6 (CANB 906471.1) | AU: NSW; Central Tablelands; Mount Canobolas State Conservation Area |
| *Dipodium* aff. *roseum* 3 | HTCG 0831 | C. Bower ORG7818 WP 7 (CANB 906471.1) | AU: NSW; Central Tablelands; Mount Canobolas State Conservation Area |
| *Dipodium* aff. *roseum* 4 | HTCG 0832 | C. Bower ORG7818 WP 9,10,11 (CANB 906471.1) | AU: NSW; Central Tablelands; Mount Canobolas State Conservation Area |
| *Dipodium* aff. *stenocheilum* | HTCG 1691 | D.L. Jones 8968 (CBG 9220253.1) | AU: QLD; Cook; Mount Elliot |
| *Dipodium* *ammolithum* | HTCG 1372 | M.D. Barrett 4910A (PERTH) | AU: WA; North Kimberley, Theda Station |
| *Dipodium* *atropurpureum* 1 | HTCG 0760 | W.M. Dowling DC 1717 (CANB 924629.1) | AU: NSW; Northern Tablelands; Barrington Tops State Forest |
| *Dipodium* *atropurpureum* 2 | HTCG 1679 | M.A. Clements 4426 (CBG 8605570.1) | AU: NSW; Northern Tablelands; New England Highway to Armidale |
| *Dipodium* *basalticum* | HTCG 1693 | D.E. Murfet 4837 (CANB 662327.1) | AU: NT; Darwin and Gulf; near Nhulunbuy |
| *Dipodium* *campanulatum* 1 | HTCG 1680 | K. Alcock DLJ5622 (CBG 9004646.2) | AU: SA; South-east; Naracoorte |
| *Dipodium* *campanulatum* 2 | HTCG 1681 | D.E. Murfet 1930b (CANB 677107.2) | AU: SA; South-east; Penola Conservation Park |
| *Dipodium* *elegantulum* | HTCG 1682 | L. Lawler 8 (CBG 8605836.1) | AU: QLD; Cook; near Mareeba |
| *Dipodium* *ensifolium* | HTCG 1343 | D.M. Crayn 1581 (CNS 145658.1) | AU: QLD; Cook; record is Queensland sensitive |
| *Dipodium* *hamiltonianum* | HTCG 1683 | D.L. Jones & P.D. Jones s.n. (CANB) | AU: QLD; Moreton; Currimundi |
| *Dipodium* *interaneum* | HTCG 0181 | J. Egan ORG7745 (CANB) | AU: ACT; Canberra; Birrigai |
| *Dipodium* *pandanum* 1 | CNS_G01262 | B. Gray 8233 (CANB 572368.2) | AU: QLD; Kennedy North; near Coen; record is Queensland sensitive |
| *Dipodium pandanum* 2 | HTCG 1694 | M. Jacobs 8984 (CANB 576763.1) | PG: Mount Bosavi |
| *Dipodium pardalinum* 1 | HTCG 1684 | D.L. Jones 12834 (CBG 9603749.1) | AU: Vic; Victorian Volcanic Plain; Heathmere |
| *Dipodium pardalinum* 2 | HTCG 1685 | D.L. Jones 12830 (CBG 9603745.1) | AU: Vic; Victorian Volcanic Plain; Heathmere |
| *Dipodium pulchellum* | HTCG 1686 | D.L. Jones s.n. (CANB) | AU: QLD; Moreton; Green Mountains |
| *Dipodium punctatum* | HTCG 0827 | C. Bower ORG7816 (CANB 906469.1) | AU: NSW; Central Tablelands; Black Salee Reserve |
| *Dipodium roseum* 1 | HTCG 1687 | C. Houston ORG3859 (CANB 656733.1) | AU: SA; Lofty South; Wotton Scrub |
| *Dipodium roseum* 2 | HTCG 1688 | C. Houston ORG3859 (CANB 656733.2) | AU: SA; Lofty South; Wotton Scrub |
| *Dipodium stenocheilum* 1 | HTCG 1689 | M.A. Clements 1189 (CBG 7801007.1) | AU: NT; Darwin and Gulf; Elcho Island |
| *Dipodium stenocheilum* 2 | HTCG 1690 | D.E. Murfet 3018 (CANB 619696.1) | AU: NT; Darwin and Gulf; Livingston |
| *Dipodium variegatum* | HTCG 1692 | D.L. Jones 1280 (CANB 665182.1) | AU: QLD; Moreton; Beenleigh |
| Outgroup |  |  |  |
| *Acriopsis emarginata* | CNS_G00305 | C.D. Kilgour 634A (CNS 135324.1) | AU: QLD, Cook, Daintree National Park |
| *Cymbidium canaliculatum* | CNS_G00165 | K.R. McDonald, 11722 (BRI AQ0831415) | AU: QLD, Cook, Mungkan Kandju National Park |
| *Eulophia bicallosa* | HTCG 1696 | I. Morris (DLJ 4579) (CBG 8913381.1) | AU: NT; Darwin and Gulf; Howard Springs |
| *Eulophia graminea* | CNS_G02766 | C.P. Brock 311 (CANB 596921.1) | AU: NT; Darwin |
| *Eulophia nuda* | HTCG 1697 | R. Crane 1072 (CANB) | cult. ex AU: QLD; Moreton; Caloundra |
| *Geodorum densiflorum* | CNS_G01890 | K. Schulte 254B (CNS 146066.1) | AU: QLD; Cairns region |
| *Oeceoclades pelorica* | HTCG 1695 | J. Taylor s.n. (CBG 7905124.1) | cult. ex AU: QLD; Cook; Iron Range |

### 2.2 DNA extraction, library preparation, and sequencing

Standard plant DNA extractions were carried out from 5-20 mg of silica dried plant tissue from field collections or herbarium material (**Table 1**) at the National Research Collections Australia (NRCA, CSIRO) in Canberra. The Invisorb DNA Plant HTS96 kit (Stratec, Birkenfeld, Germany) was used following the manufacturer’s protocol, with a final elution of 60 ml.

DNA of *Dipodium* samples (**Table 1**) was sonicated to an average target length of ca. 200 bp using a LE220 sonicator (Covaris, Bankstown, Australia). After sonication, DNA length and concentration were quantified on Fragment Analyzer (Agilent Technologies, California, USA) using the Agilent high-sensitivity genomic DNA kit.

DNA libraries were prepared using the QiaSeq UltraLow Input library kit (Qiagen, Germantown, Australia) using custom dual-indexed adapters. Final libraries were size-selected on Fragment Analyzer using the high-sensitivity Genomic Fragment Analyzer Kit (Agilent, Santa Clara, USA), quantified using the Fluoroskan plate fluorometer (Thermo Fisher Massachusetts, USA) and the Quant-iT HS dsDNA kit (Invitrogen, California, USA) following the manufacturer’s instructions. Samples were pooled equimolarly and sequenced using 150 bp paired end reads on a NovaSeq S1 flowcell (Illumina, California, USA) at the Biomolecular Resource Facility within the John Curtin School of Medical Research, Australian National University (Canberra, Australia).

### 2.3 Data processing and whole plastid genome assembly

We carried out both *de novo* and reference-guided assemblies for the *Dipodium* data set. Trimming and assembly of *de novo* contigs were carried out as described in Nargar et al. (2022). Briefly, raw sequences were trimmed applying a Phred score > 20 using Trimmomatic 0.39 (Bolger et al., 2014), and deduplicated using ‘clumpify’ from BBtools 38.9 (Bushnell, 2014). Read pairs were then assembled using SPAdes 3.15 (Bankevich et al., 2012). Plastid databases were extracted from NCBI’s Nucleotide Entrez database using Entrez Programming Utilities (2008) using taxonomic, keyword, and sequence length constraints. Contigs were identified as derived from plastid source using blastn against these databases. Genes within plastid contigs were identified by homology using BLAST (Altschul et al., 1990) and BLASTx (RRID:SCR_001653) against genes extracted from annotations of the reference sequence sets extracted from nuccore.

Reference-guided assemblies were performed with paired, merged reads and the recently published and closely related plastome of *Dipodium roseum* D.L.Jones and M.A.Clem. (MN200386, Kim et al., 2020). The related orchid *Masdevallia coccinea* Linden ex Lindl. (KP205432, Kim et al., 2015) was included as an additional reference sequence to ensure that regions which already showed degradation in some plastid genes in the plastome of *D. roseum* (e.g., all *ndh* genes) (MN200386, Kim et al., 2020) and which may still be present in other *Dipodium* species could be assembled as the plastome of *M. coccinea* has a full set of functional plastid genes (Kim et al., 2015).

Reference-guided assemblies were carried out using the plugin ‘map to reference’ in Geneious Prime (Version 2022.0.2, Biomatters Ltd, www.geneious.com) with default settings. To obtain complete plastome assemblies, consensus sequences for each sample were extracted (threshold 60%, reading depth > 10), aligned using MAFFT v7.388 (Katoh and Standley 2013) in Geneious, manually checked and compared. Reference-guided assemblies were visually inspected and in cases of misassembled regions due to potential mismatches between the sample and the reference de novo assemblies were consulted, and were quality allowed the region extracted from the de novo assembly. The prediction and finding of gene annotations for complete plastome assemblies were performed with the Geneious plugin ‘predict annotation’ (similarity: 90% and best match with *D. roseum* (MN200386)). Open reading frames (ORFs) were manually checked and verified by identifying the start and stop codons. In cases of remaining ambiguities, BLAST searches were conducted for reading-frame verification (Altschul et al. 1990; National Center for Biotechnology Information; Available from: https://blast.ncbi.nlm.nih.gov/Blast.cgi [cited: 08 Sept 2023]). The inverted repeat (IR) boundaries were identified using the ‘repeat finder’ plugin in Geneious with default settings. In total, 24 complete *Dipodium* plastomes were assembled in this study. The graphical representation of each plastome and divergent regions with annotations were created in OrganellarGenomeDRAW (OGDRAW, version 1.3.1, Greiner et al., 2019).

### 2.4 Phylogenetic analyses

To elucidate phylogenetic relationships within *Dipodium* and to assess the phylogenetic position of *Dipodium* within Cymbidieae we performed a phylogenetic analysis with DNA sequences of 33 newly sequenced plastomes from this study (**Table 1**) and an extended outgroup sampling for 115 samples from published plastid data (Supplementary Material 1). Coding regions of respective genes of 33 samples were extracted with the ‘extract’ function in Geneious Prime. Where mutations had led to frame shifts with internal stop codons, the affected sequences were excluded from phylogenetic analyses.

Each extracted coding region of in total 68 plastid loci from 33 samples (including the intron regions) and from 115 published plastomes (excluding intron regions) were aligned using MAFFT (v7.388; Katoh et al., 2002; Katoh and Standley 2013) Geneious prime plugin with default settings, checked manually and subsequently concatenated to an alignment of 69,335 bp (Supplementary Material 2).

Maximum likelihood analysis of the plastid dataset (148 samples) with best-fit models GTR+I+I+F+R4 was performed using IQ-TREE ver. 2.2.0 (Nguyen et al., 2015; Kalyaanamoorthy et al., 2017; Minh et al., 2020). Branch support was obtained with Shimodaira-Hasegawa-like approximate Likelihood Ratio Test (SH-aLRT; Guindon et al., 2010) and the ultrafast bootstrap (ufboot2; Hoang et al., 2018) as implemented in the IQ-TREE software. The tree topology was visualised using the software Figtree (ver. 1.4.4.; http://tree.bio.ed.ac.uk/software/figtree/).

### 2.5 Divergence-time analysis

For divergence-time estimations of *Dipodium*, the alignments were reduced to the 30 most parsimony informative loci due to computational limitations. The 30 plastid loci were selected based on their most parsimony informative (Pi) sites estimated with MEGA (Molecular Evolutionary genetics Analysis; ver. 11.0.11, Tamura et al., 2021) and presence of loci across the dataset (Supplementary Material 2). For taxa represented by more than one sample, duplicates were removed from alignments as recommended for divergence time estimation. Alignments of 30 plastid loci from 134 taxa were concatenated yielding a total alignment length of 27,934 bp using MAFFT (v7.388; Katoh et al., 2002; Katoh and Standley 2013) implemented in Geneious Prime (Supplementary Material 2). Absolute node ages and phylogenetic relationships were jointly estimated in BEAST (ver. 2.7.4; Bouckaert et al., 2019, Bouckaert et al. 2014) applying the best fit partition scheme and substitution model as determined by IQ-TREE’s ModelFinder (GTR+F+I+I+R4). Four different models were tested: a Bayesian optimised relaxed and a strict molecular clock with uncorrelated lognormal rates with each a Yule and a Birth-death tree prior on the speciation process (Douglas et al., 2021; Gernhard et al., 2008; Zuckerkandl and Pauling, 1965; Yule, 1925). Trees were calibrated with four secondary calibration points based on Zhang et al. (2023). A normal distribution with an offset value of 101.52 Ma and a standard deviation (SD) of 2.2 was assigned as crown age of Orchidaceae. The priors for the three other calibration points were set with a normal distribution and the means of stem ages for Vanilloideae (offset value = 93.48 Ma, SD = 2.7), Cypripedioideae (offset value = 89.14 Ma, SD = 2.71) and Orchidoideae (offset value = 77.74 Ma, SD = 2.0). For each clock model, 10 parallel BEAST analyses with each 30 million generations and a sampling frequency of every 10,000 generations were carried out. The run parameters were examined in TRACER (ver. 1.7.2; Rambaut et al., 2018) and the effective sample sizes (ESSs) of > 200 for all parameters and the burn-in were assessed. The runs were combined in LogCombiner (Drummond and Rambaut 2007) with a burn-in of 10% and subsequently used to generate a maximum-clade-credibility chronogram with mean node heights in TreeAnnotator (Drummond and Rambaut 2007). To determine the best fitting clock model and speciation models for the data set, a model comparison using the AICM (Akaike Information Criterion by MCMC) was performed with BEAST v.2.6.2 and evaluated with the AIC model selection criterion of Fabozzi et al. (2014).

### 2.6 Plastid genome evolution

#### 2.6.1 Structural variation in *Dipodium* plastomes

To examine structural variation among the plastomes of *Dipodium,* whole plastome alignments were generated using MAFFT (v7.388; Katoh et al., 2002; Katoh and Standley 2013) implemented in Geneious Prime with full annotations. Alignments were manually checked, in cases of divergent regions e.g., the operon region of *ndh*C, *ndh*K, and *ndh*J genes or junctions between the large single copy (LSC)/ inverted repeat B (IRB)/ small single copy (SSC)/ inverted repeat A (IRA) regions, and respective regions (including annotations) were extracted in Geneious Prime, separately aligned, proofread, and subsequently visualised using OGDRAW (ver. 1.3.1, Greiner et al., 2019).

#### 2.6.2 Functional genes, pseudogenes, and physical gene loss

To classify the level of degradation of plastid genes in *Dipodium*, we used the following categories: (1) ***functional*** - the reading frame was intact and less than 10% of the open reading frame was disrupted by small indels; (2) ***moderately pseudogenised*** - less than 10% of the open reading frame was disrupted by internal stop codons or indels causing non-triplet frame shifts; (3) ***severely pseudogenised*** - more than 10% of the open reading frame was disrupted by either internal stop codons, large deletions (> 10%), and non-triplet frame shifts (based on Barrett et al., 2019), or (4) ***lost*** - the gene was not identified in the annotation process of the *de-novo* assembly (e.g., Joyce et al., 2018) and/or was not detectable within the reference-guided assembly. A gene was considered as not detectable within the reference-guided mapping process if at least 70% of the gene sequence could not be identified for calculation of the consensus sequence within the Geneious mapping process. The coded matrix of gene degradation was plotted against the maximum likelihood phylogenetic tree of *Dipodium*.

## 3 Results

### 3.1 Phylogenetic placement of *Dipodium* in tribe Cymbidieae and infrageneric relationships within the genus

The maximum likelihood analysis based on 68 plastid loci and 148 samples yielded highly resolved and well-supported tree topologies for the phylogenomic relationships within Orchidaceae (Supplementary Material 3). Within Epidendroideae, Cymbidiinae was monophyletic and sister to all other Cymbidieae including Dipodiinae (SH-aLRT/UFboot 100/100; **Figure 2**). *Dipodium* was retrieved as next diverging lineage within Cymbidieae and monophyletic with maximum support values (SH-aLRT/UFboot 100/100; **Figure 2**).

**Figure 2:**
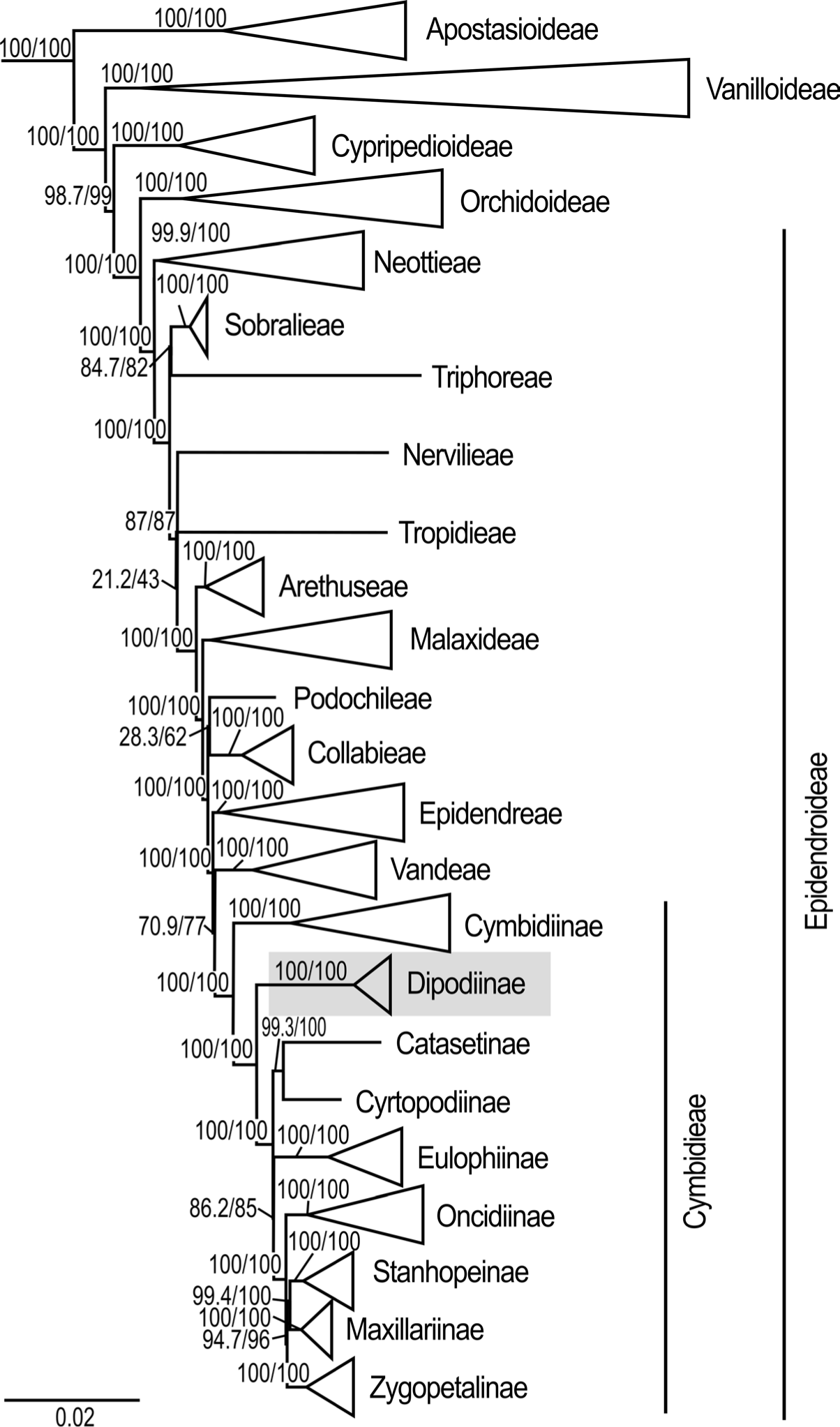
Phylogenetic relationships among major orchid lineages and placement of subtribe *Dipodiinae* in Cymbidieae. Maximum likelihood tree of 148 taxa based on 68 plastid loci. Support values are shown above each branch, SHaLRT followed by UFBoot values. Scale bar represents branch length, along which 0.02 per-site substitutions are expected. Detailed phylogeny provided in Supplementary Material 3.

Within *Dipodium*, section *Leopardanthus* was placed as sister group to section *Dipodium* with maximum support values (SH-aLRT/UFboot 100/100; **Figure 3**).

**Figure 3:**
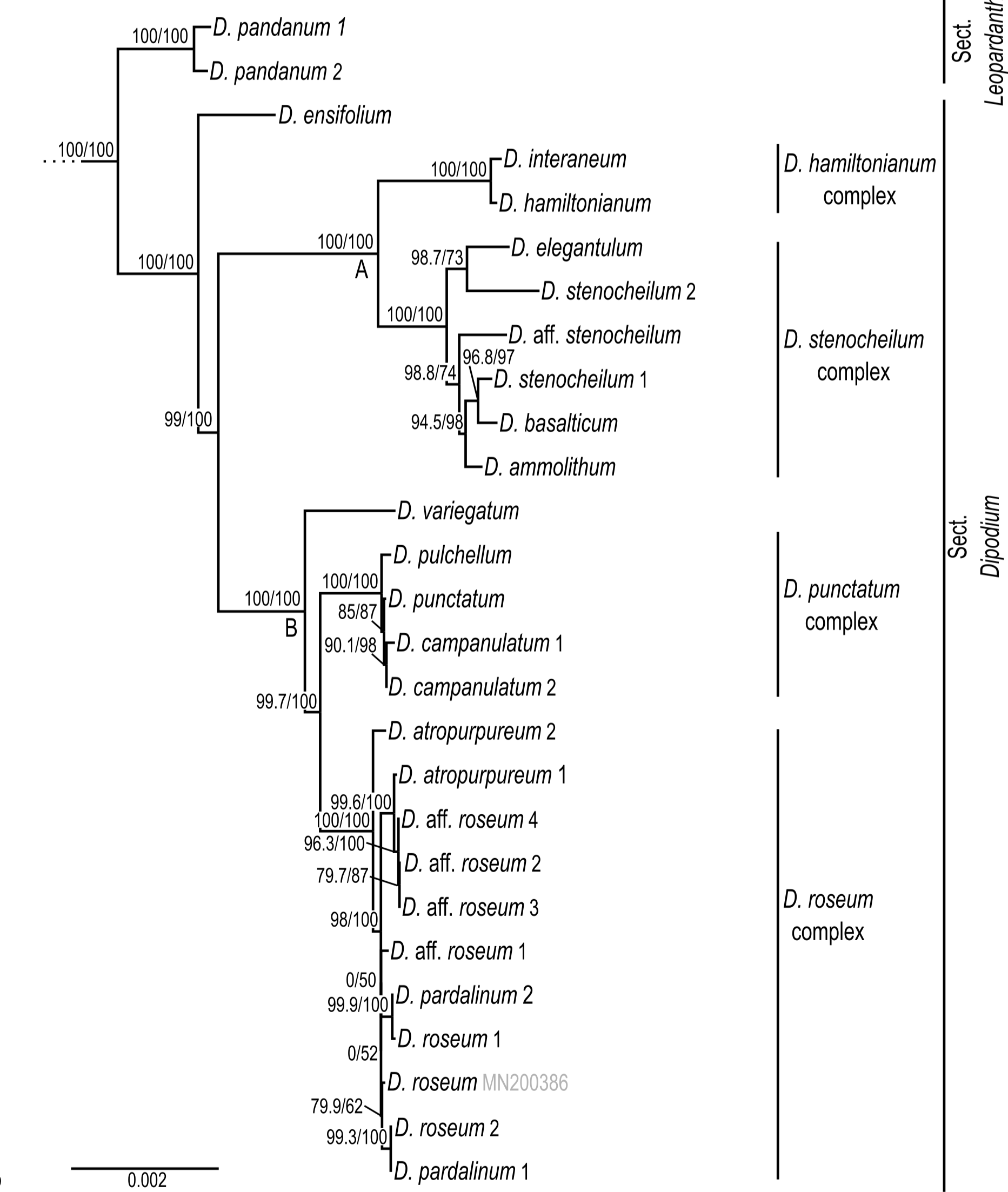
Phylogenetic relationships in *Dipodium.* Maximum likelihood tree based on 68 plastid loci and 148 taxa (outgroups not shown). Support values are given above each branch, SHaLRT is followed by UFBoot values. Scale bar represents branch length, along which 0.002 per-site substitutions are expected.

Section *Dipodium* was resolved as monophyletic and divided into six highly supported lineages. The leafy species *D. ensifolium* was placed as sister to all leafless species of the section (SH-aLRT/UFboot 100/100; **Figure 3**). Next, sect. *Dipodium* split into two main clades, A and B (SH-aLRT/UFboot 99/100; **Figure 3**). Clade A split into two lineages, the *Dipodium hamiltonianum* complex and the *Dipodium stenocheilum* complex, receiving maximum nodal support (SH-aLRT/UFboot 100/100; **Figure 3**). The *D. hamiltonianum* complex comprised the two species *D. hamiltonianum* and *D. interaneum.* The *D. stenocheilum* complex included *D. ammolithum, D. basalticum, D. elegantulum, D. stenocheilum*, and *D.* aff. *stenocheilum. Dipodium stenocheilum* was retrieved as non-monophyletic. (**Figure 3**).

Clade B resolved *D. variegatum* as sister to the remaining species of the clade (SH-aLRT/UFboot 100/100). The remainder split into the *D. punctatum* complex and the *D. roseum* complex (SH-aLRT/UFboot 99.7/100; **Figure 3**). The *D. punctatum* complex comprised three species, *D. campanulatum, D. pulchellum,* and *D. punctatum*. Phylogenetic divergence between these three species was shallow and support for interspecific relationships within the complex low. The *D. roseum* complex comprised four taxa, namely *D. atropurpureum, D. pardalinum, D. roseum,* and *D.* aff. *roseum*. Resolution and support for interspecific relationships within the *D. roseum* complex was low overall.

### 3.2 Divergence-time estimations

Absolute times of divergence under strict and optimised relaxed clocks for Orchidaceae based on 30 plastid loci and 134 taxa showed similar results. Strict clock models consistently yielded slightly older age estimates than the analyses based on the relaxed clock models (Supplementary Material 4). Model comparison using AICM (Fabozzi et al., 2014) identified the relaxed clock model under the birth-death speciation model as the best fit models for the dataset (Supplementary Material 4).

The Bayesian relaxed clock tree topology and the maximum likelihood phylogeny agreed overall in major relationships within Orchidaceae and the placement of species within *Dipodium*. Epidendroideae were estimated to have emerged ca. 77.7 Ma (HDP: 74.2–81.5) with the stem age of subtribe Cymbidieae placed in the Eocene, ca. 42.2 Ma (HDP: 34.3–50.1) (Supplementary Material 4 and 5). The stem age of subtribe Cymbidiinae, the first diverging lineage in Cymbidieae, was placed in the late Eocene, ca. 38.0 Ma (HDP: 30.7–45.7) (**Figure 4**). Stem diversification of Dipodiinae was estimated to have commenced ca. 33.3 Ma (HDP: 26.4–40.6) in the early Oligocene (**Figure 4**). Crown diversification of Dipodiinae was estimated to have commenced much later, in the late Miocene with sections *Dipodium* and *Leopardanthus* diverging ca. 11.3 Ma (HDP: 6.8–16.2) (**Figure 4**). The crown age of section *Dipodium* was estimated to be ca. 8.1 Ma (HDP: 5.2–11.6) in the late Miocene with the divergence of the leafy species, *D. ensifolium*, from the remainder of section *Dipodium* (**Figure 4**). The crown age of the remainder of the section, i.e., all leafless species, was estimated to ca. 7.3 Ma (HDP: 4.4–10.4) (**Figure 4**). Within this leafless clade, two subclades each containing two species complexes were resolved. The crown age of the clade comprising the *D. hamiltonianum* complex and the *D. stenocheilum* complex was estimated to ca. 4.3 Ma (HDP:2.5–6.4) in the early Pliocene (**Figure 4**) which is congruent with estimations of the crown age of clade B (comprising *D. variegatum* and the two complexes *D. punctatum* and *D. roseum*) (**Figure 4**). The *D. stenocheilum* complex had a crown age of ca. 2.4 Ma (HDP: 1.3– 3.6) in the early Pleistocene. The three remaining complexes had crown ages estimated to the mid Pleistocene (*D. hamiltonianum* complex: ca. 0.7 Ma, HDP: 0.1–1.4; *D. punctatum complex*: ca. 0.6 Ma, HDP: 0.2–1.0, and *D. roseum* complex: ca. 0.7 Ma, HDP: 0.3–1.3) (**Figure 4**).

**Figure 4:**
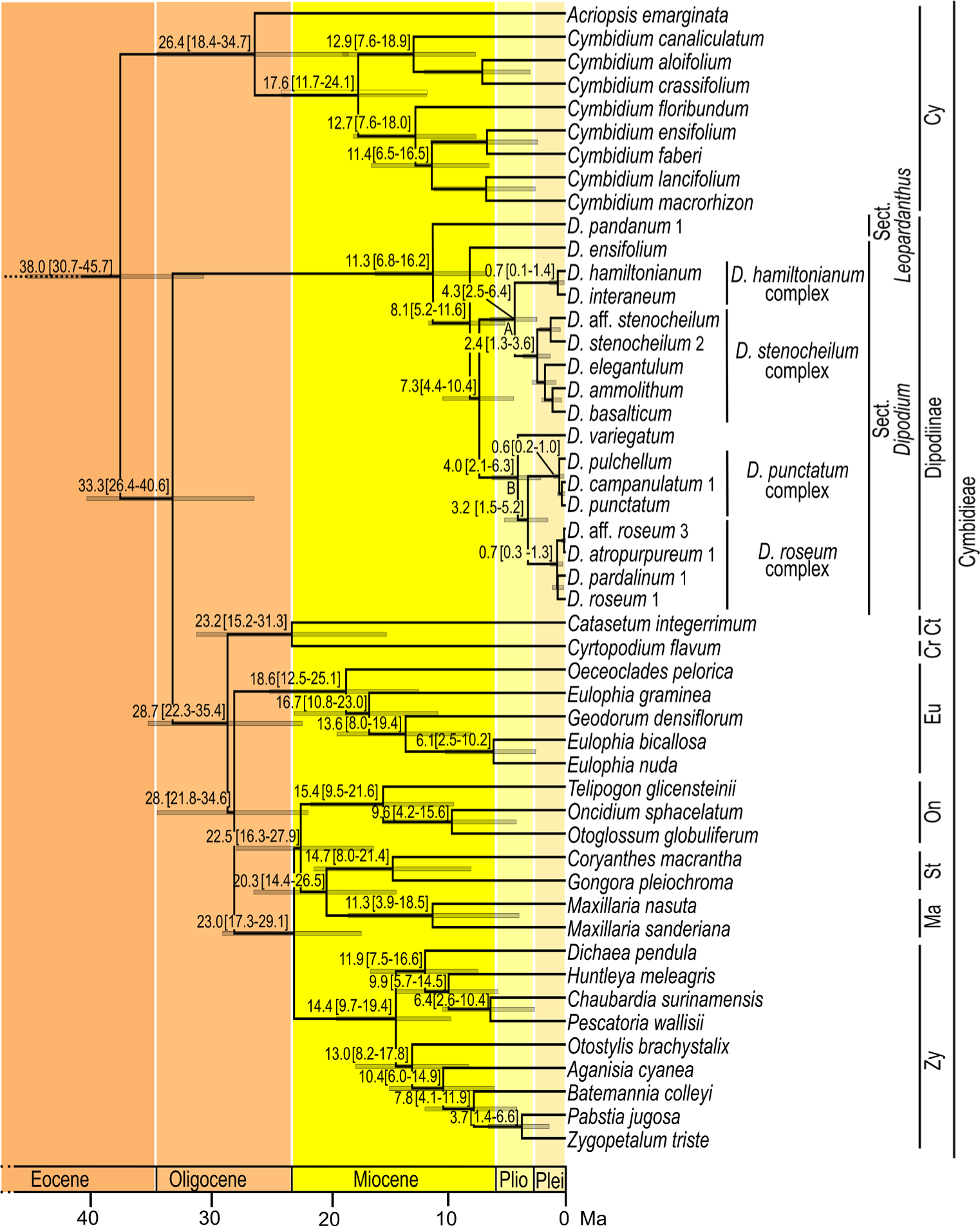
Chronogram of Cymbidieae. Maximum-clade-credibility tree from Bayesian divergence-time estimation in BEAST2 based on 30 plastid loci and an optimised lognormal molecular clock model under the birth-death prior (outgroups not shown). Divergence times (million years ago) are shown at each node, together with 95% highest posterior density (HDP) values indicated by grey bars and values in parentheses. A and B refers to the two main lineages within sect. *Dipodium*. Cy: Cymbidiinae, Ct: Catasetinae, Cr: Cyrtopodiinae, Eu: Eulophiinae, On: Oncidiinae, St: Stanhopeinae, Ma: Maxillariinae, Zy: Zygopetalinae, Plio: Pliocene, Plei: Pleistocene. Outgroups to Cymbidieae not shown. Detailed chronogram provided in Supplementary Material 5.

### 3.3 Characterisation of *Dipodium* plastomes

Complete plastome assemblies and annotations were successfully carried out for 24 *Dipodium* samples, representing all Australian species of section *Dipodium* including two recently discovered species of section *Dipodium* (*D. ammolithum* and *D. basalticum*), two putatively new species of section *Dipodium* (*D.* aff. *roseum*, *D.* aff. *stenocheilum*) and one species of section *Leopardanthus* (*D. pandanum*) (**Table 2**). Plastome assemblies for *D. pandanum* 2 and *D*. aff. *stenocheilum* showed an insufficient mean coverage (<30) for non-coding regions which caused unsolved gaps and ambiguous bases which could not be reliably resolved. The number of paired-end, trimmed reads for the successfully assembled complete plastomes ranged from 332,604 (*D. pandanum* 1) to 27,999,734 (*D. pardalinum* 2) and the mean coverage ranged from 31x to 627x (Supplementary Material 6).

**Table 2.** Comparison of plastome features in *Dipodium.*

| Sample | Plastome<br>Length<br>(bp) | SSC<br>length<br>(bp) | IRA/B<br>length<br>(bp) | LSC<br>length<br>(bp) | GC<br>content | Total<br>CDS<br>(unique<br>CDS) | Total<br>tRNA<br>(unique<br>tRNA) | Total<br>rRNA<br>(unique<br>rRNA) | Total<br>pseudo-<br>genes | Total<br>lost<br>genes | Total<br>functional<br>genes |
| --- | --- | --- | --- | --- | --- | --- | --- | --- | --- | --- | --- |
| *D. pandanum* 1 | 146,204 | 13,849 | 24,762 | 82,831 | 37.0% | 74 (68) | 38 (30) | 8 (4) | 9 | 3 | 120 |
| *D. ensifolium* | 150,084 | 16,756 | 25,497 | 82,334 | 36.9% | 74 (68) | 38 (30) | 8 (4) | 10 | 3 | 120 |
| *D. hamiltonianum* complex |  |  |  |  |  |  |  |  |  |  |  |
| *D. hamiltonianum* | 145,902 | 14,384 | 24,929 | 81,660 | 37.1% | 74 (68) | 39 (31) | 8 (4) | 10 | 3 | 121 |
| *D. interaneum* | 146,497 | 14,635 | 24,951 | 81,960 | 37.0% | 74 (68) | 39 (31) | 8 (4) | 10 | 3 | 121 |
| *D. stenocheilum* complex |  |  |  |  |  |  |  |  |  |  |  |
| *D. elegantulum* | 144,865 | 14,003 | 24,606 | 81,650 | 36.9% | 74 (68) | 39 (31) | 8 (4) | 9 | 4 | 121 |
| *D. stenocheilum* 2 | 145,589 | 13,821 | 25,127 | 81,514 | 37.0% | 74 (68) | 39 (31) | 8 (4) | 9 | 4 | 121 |
| *D. stenocheilum* 1 | 144,751 | 12,670 | 25,009 | 82,063 | 36.9% | 74 (68) | 38 (30) | 8 (4) | 9 | 4 | 120 |
| *D. basalticum* | 148,478 | 15,238 | 25,640 | 81,960 | 37.0% | 74 (68) | 38 (30) | 8 (4) | 10 | 3 | 120 |
| *D. ammolithum* | 147,842 | 14,697 | 25,600 | 81,946 | 37.0% | 74 (68) | 39 (31) | 8 (4) | 9 | 4 | 121 |
| *D. variegatum* | 142,949 | 12,039 | 24,436 | 82,038 | 37.0% | 74 (68) | 38 (30) | 8 (4) | 6 | 7 | 120 |
| *D. punctatum* complex |  |  |  |  |  |  |  |  |  |  |  |
| *D. pulchellum* | 151,425 | 15,735 | 26,369 | 82,952 | 36.9% | 74 (68) | 38 (30) | 8 (4) | 11 | 2 | 120 |
| *D. punctatum* | 151,181 | 15,737 | 26,136 | 83,172 | 37.0% | 74 (68) | 38 (30) | 8 (4) | 11 | 2 | 120 |
| *D. campanulatum* 1 | 146,390 | 13,602 | 25,284 | 82,220 | 36.9% | 74 (68) | 37 (29) | 8 (4) | 12 | 2 | 119 |
| *D. campanulatum* 2 | 149,050 | 14,266 | 25,902 | 82,980 | 37.0% | 74 (68) | 38 (30) | 8 (4) | 11 | 2 | 120 |
| *D. roseum* complex |  |  |  |  |  |  |  |  |  |  |  |
| *D. atropurpureum* 2 | 149,390 | 15,509 | 25,909 | 82,063 | 36.9% | 74 (68) | 38 (30) | 8 (4) | 10 | 3 | 120 |
| *D. atropurpureum* 1 | 150,481 | 15,633 | 26,399 | 82,050 | 36.9% | 74 (68) | 38 (30) | 8 (4) | 9 | 4 | 120 |
| *D. aff. roseum* 4 | 152,282 | 16,426 | 26,630 | 82,596 | 36.9% | 73 (67) | 38 (30) | 8 (4) | 11 | 3 | 119 |
| *D. aff. roseum* 2 | 150,462 | 15,514 | 26,388 | 82,172 | 36.9% | 74 (68) | 38 (30) | 8 (4) | 9 | 4 | 120 |
| *D. aff. roseum* 3 | 152,956 | 16,571 | 26,817 | 82,751 | 36.9% | 74 (68) | 38 (30) | 8 (4) | 10 | 3 | 120 |
| *D. aff. roseum* 1 | 151,791 | 16,362 | 26,424 | 82,581 | 36.9% | 74 (68) | 38 (30) | 8 (4) | 10 | 3 | 120 |
| *D. pardalinum* 2 | 151,659 | 16,276 | 26,580 | 82,223 | 36.9% | 74 (68) | 38 (30) | 8 (4) | 10 | 3 | 120 |
| *D. pardalinum* 1 | 148,174 | 15,283 | 25,494 | 81,903 | 36.8% | 74 (68) | 38 (30) | 8 (4) | 9 | 4 | 120 |
| *D. roseum* 1 | 150,857 | 15,848 | 26,521 | 81,967 | 36.9% | 74 (68) | 38 (30) | 8 (4) | 9 | 4 | 120 |
| *D. roseum* 2 | 147,730 | 14,192 | 25,819 | 81,900 | 36.8% | 74 (68) | 38 (30) | 8 (4) | 9 | 4 | 120 |

#### 3.3.1 Plastome features and structural variations within *Dipodium* plastomes

Plastome sizes of *Dipodium* ranged from 142,949 bp (*D. variegatum*) to 152,956 bp (*D*. aff. *roseum* 3) (**Table 2**, **Figure 5**, Supplementary Material 7). The largest average plastome size (150,578 bp) was found in the *D. roseum* complex, closely followed by the leafy *D. ensifolium* (150,084 bp), and the *D. punctatum* complex (149,512 bp). Plastome sizes within the *D. stenocheilum* complex were markedly lower with an average size of 146,305 bp. Similarly small plastomes were also found in *D. hamiltonianum* (145,902 bp), *D. interaneum* (146,497 bp) and the leafy climber *D. pandanum* 1 (sect. *Leopardanthus*) (146,204 bp) (**Table 2**). *Dipodium* plastomes possess the typical quadripartite structure of angiosperms, with the SSC region ranging from 12,039 bp (*D. variegatum*) to 16,756 bp (*D. ensifolium*), the LSC region ranging from 81,514 bp (*D. stenocheilum* 2) to 83,172 bp (*D. punctatum*), and the pair of IRs ranging from 24,436 bp (*D. variegatum*) to 26,817 bp (*D*. aff. *roseum* 3) (**Table 2**).

**Figure 5:**
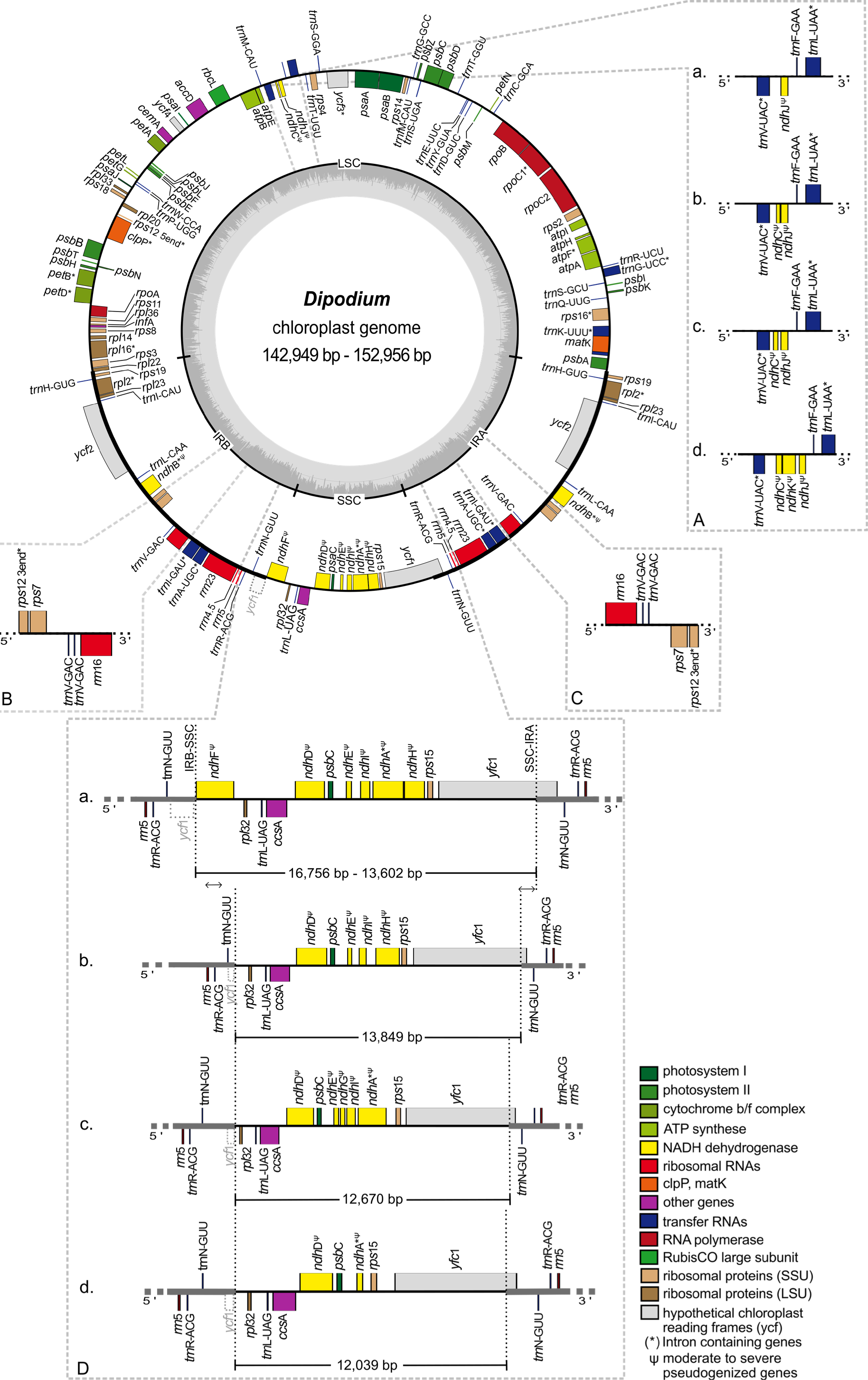
Plastome map and boundary shifts in *Dipodium*. The plastome of *D. atropurpureum* 2 is illustrated as representative and shown as a circular gene map with the smallest and the largest *Dipodium* plastome of this study. Genes outside the circle are transcribed in a clockwise direction, those inside the circle are transcribed in a counterclockwise direction. The dark grey inner circle corresponds to the G/C content, and the lighter grey to the A/C content. The major distinct regions of complete *Dipodium* plastomes are compared in each detailed enlarged box (A-D). (A) Note that each representative block (a-d) has pseudogenised or lost either *ndh*J, *ndh*K or *ndh*C genes. (B, C). Duplication of *trn*V-GAC in the Inverted Repeat regions of *D. interaneum* (IRB), *D. hamiltonianum* (IRA), *D. elegantulum* (IRB), *D. stenocheilum* 2 (IRA), *D. ammolithum* (IRA). (D) Each block (a. as representative *D. roseum* 2; b. *D. pandanum* 1; c. *D. stenocheilum* 1; d. *D. variegatum*) shows differences in the length (bp) of the SSC region caused through loss or pseudogenization of either *ndhF*, *ndhD*, *ndhE*, *ndh*G, *ndh*I, *ndh*A or *ndh*H, note the boundary shift of the IRs/SSC region caused through the loss/ pseudogenisation of *ndh*F and the inclusion of the functional *ycf*1 and the *ycf1*-fragment (grey, dashed line) into the IRs. SSC: Small Single Copy; LSC: Large Single Copy: IRA/B: Inverted Repeat A/B.

Total mean GC content of *Dipodium* plastomes was 36.9%, ranging between 36.8% (*D. roseum* 2 and *D. pardalinum* 1) and 37.1% (*D. hamiltonianum*) (**Table 2**). Within the *D. roseum* complex the GC content was 36.8% – 36.9%, followed by the *D. punctatum* complex (36.9%), *D. stenocheilum* complex (37.0%) and the highest GC content was 37.1% and 37.0% (*D. hamiltonianum* and *D. interaneum*) (**Table 2**).

The plastid genes of each plastome were rated as functional; moderately to severely pseudogenised; or physically lost. The total number of functional genes in *Dipodium* plastomes ranged slightly from 119 to 121 including a total of 73 or 74 functional protein-coding sequence regions (CDS) (68 or 69 unique CDS), 37 to 39 functional tRNA genes (30 or 31 unique tRNA genes) and 8 rRNA genes (4 unique rRNA genes) (**Table 2**).

The IR region was largely conserved among all examined *Dipodium* plastomes. All species showed six duplicated coding regions in the IRs (i.e., *rpl*2, *rpl*23, rps7, *rps*12, *rps*19, *ycf*2) and all four rRNA genes (**Table 3**). Most plastomes showed eight duplicated tRNA genes in the IR regions with exception of the plastomes of *D. interaneum* and *D. elegantulum* which comprised a duplicated *trn*V-GAC within the IRB and the plastomes of *D. ammolithum, D. hamiltonianum,* and *D. stenocheilum* 2 which contained a duplicated *trn*V-GAC within the IRA (**Table 3**, **Figure 5**, B & C). All plastomes contained 16 functional intron-genes (i.e., *atp*F, *clp*P, *pet*B, *pet*D, *rpl*2, rpl16, *rpo*C1, *rps*12, *rps*16, *trn*A-UGC, *trn*G-UCC, *trn*I-GAU, *trn*K-UUU, *trn*L-UAA, *trn*V-UAC, *ycf*3), except for *D. pandanum* 1 which possessed two pseudogenes with introns (i.e., *ndh*A, *ndh*B) (**Table 3**, **Figure 5**). The *rps*12 gene was trans-spliced with the 5’ end located in the LSC region and 3’ end was duplicated in the IRs in all studied plastomes (**Figure 5**, Supplementary Material 7).

**Table 3.**
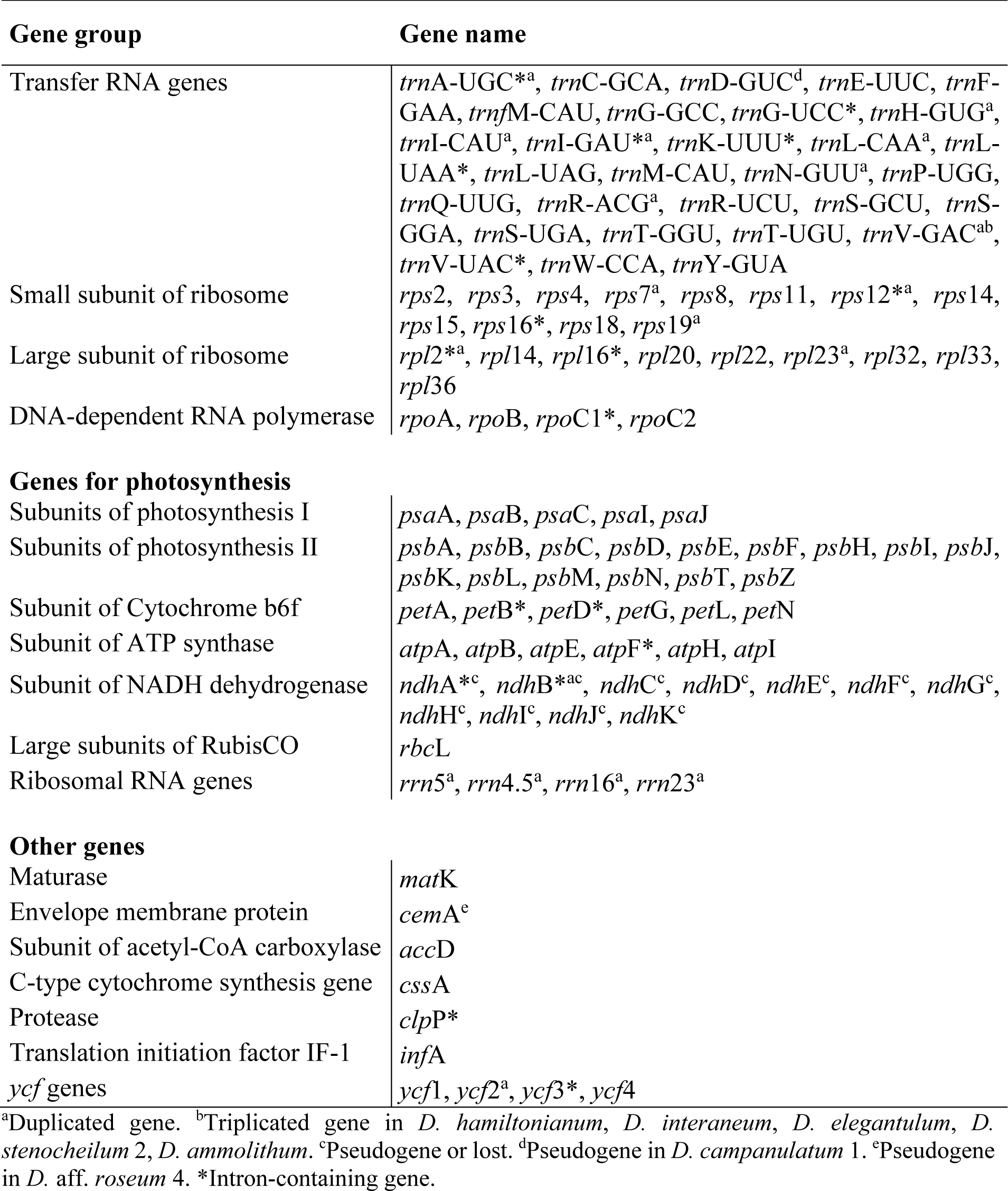
List of genes identified in the plastomes of Dipodium.

The SSC region was found to vary the most among the examined samples. All plastomes showed a contraction of the SSC with a reduction of 20-40% compared to the average size of the angiosperm SSC regions (ca. 20 kb) (Ruhlman and Jansen 2014).

Three plastomes (*D. pandanum* 1, *D. stenocheilum* 1, *D. variegatum*) lost the *ndh*F gene. This complete loss of the *ndh*F gene resulted in the *ycf*1 fragment being located in the vicinity of the *rpl*32 (**Figure 5**, D, b-d) and caused a boundary shift of the IRB/SSC region located at the 3’ end of the *ycf*1 fragment and spacer region of *rpl*32 (**Figure 5**). While all other plastomes exhibited a severely truncated *ndh*F gene but did not exhibit an IRB/ SSC boundary shift (**Figure 5**, Supplementary Material 7). The IRA/SSC junction in all examined plastomes was located within the 5’ portion of the functional *ycf*1 gene, ranging from 97 bp (*D. pandanum* 1) to 1,072 bp (*D.* aff. *roseum* 3) (**Figure 5**).

In contrast to the instability of the IR/SSC boundaries, IR/LSC boundaries were found to be relatively stable. For all studied plastomes, the LSC/IRA boundaries were located near the 3’ end of *psb*A (**Figure 5**). Variations within the LSC regions were limited to the operon which contained *ndh*C, J, K (**Figure 5**, A) and the independent pseudogenisation of *cem*A in the plastome of *D.* aff. *roseum* 4 and *trn*D-GUC in the plastome of *D. campanulatum* (**Table 3**, Supplementary Material 7).

#### 3.3.2 *ndh* gene degradation and loss in *Dipodium*

All *ndh* genes exhibited varying degrees of putative loss or pseudogenisation; not a single *ndh* gene remained functional in the examined *Dipodium* plastomes (**Table 3**, **Figure 5 & 6**). The most severe *ndh* gene loss occurred in the plastome of *D. variegatum*, with *ndh*A, *ndh*B, *ndh*D and *ndh*K severely pseudogenised and *ndh*J moderately pseudogenised.

The greatest degradation processes within *Dipodium* occurred for the *ndh*G gene, which was putatively lost in almost all plastomes, except *D. stenocheilum* 1 which retained a severely pseudogenised *ndh*G gene (**Figure 6**). This was followed by *ndh*K, which was lost in 19 out of 24 plastomes (*D. ensifolium*, D*. hamiltonianum*, *D. interaneum*, the *D. roseum* complex and the *D. stenocheilum* complex) (**Figure 6**). In the six remaining plastomes *ndh*K was conserved to different degrees. *D. punctatum*, *D. pulchellum* and *D. campanulatum* 2 retained more than 90% of the homologous bases compared to the functional *ndh*K gene of *M. coccinea* (KP205432, Kim et al., 2015) but showed severe frameshift mutations and indels which caused several internal stop codons. The plastomes with severely pseudogenised *ndh*K genes exhibited large truncations.

**Figure 6:**
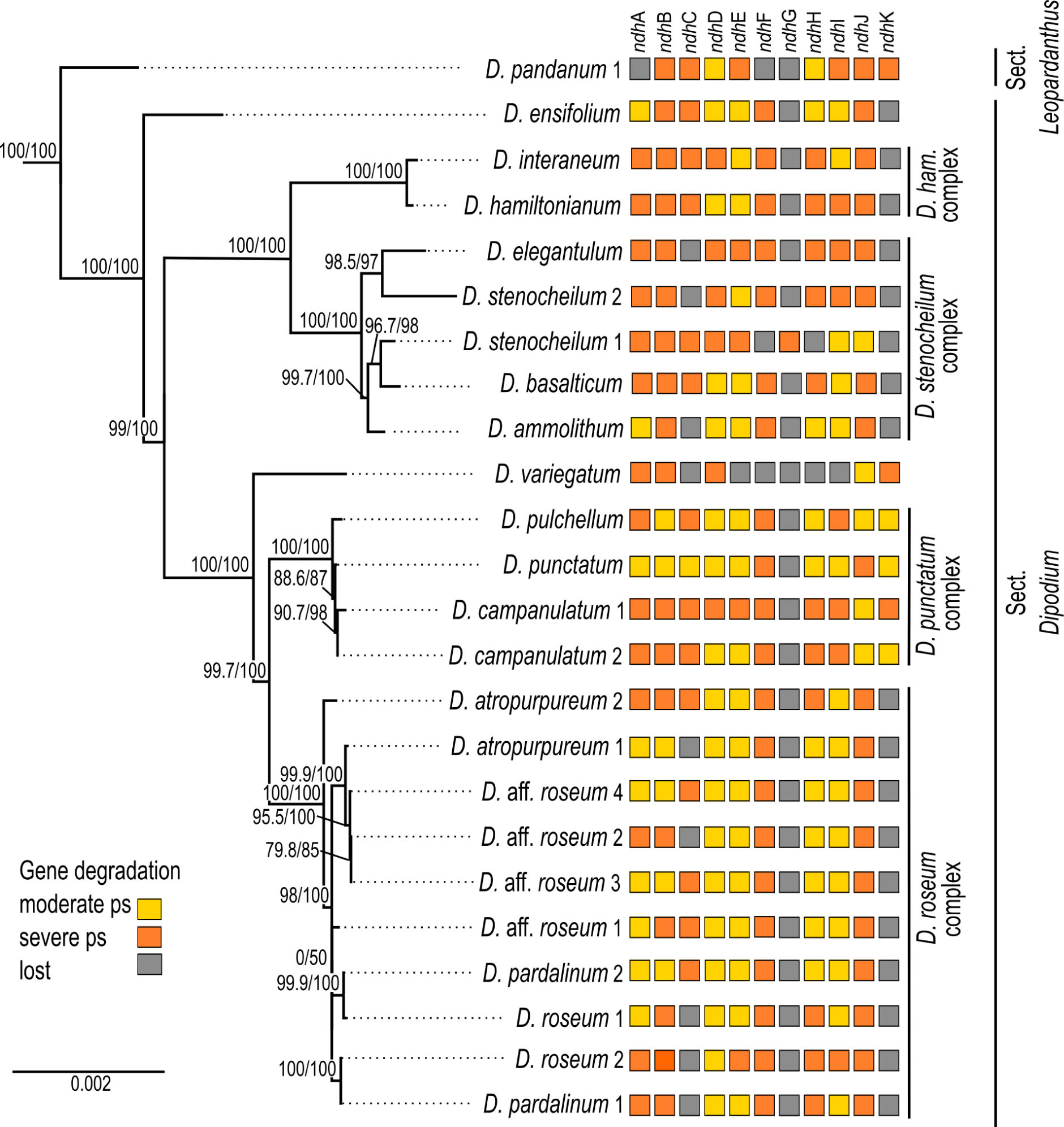
Pattern of putative *ndh* gene degradation in *Dipodium.* Gene degradation plotted against the maximum likelihood tree with focus on 24 fully assembled plastomes. (outgroups not shown). Support values (SHaLRT/ UFboot) are shown on each branch. ps = pseudogenisation; *D. ham*. = *D. hamiltonianum*

Nine *Dipodium* samples (*D. ammolithum, D. atropurpureum* 1, *D. elegantulum*, *D. pardalinum* 1, *D. roseum* 1 & 2, *D.* aff. *roseum* 2, *D. stenocheilum* 2, and *D. variegatum*) putatively lost the *ndh*C gene. In *D. punctatum ndh*C was moderately pseudogenised and in the remaining plastomes *ndh*C was severely pseudogenised (**Figure 6**). Only *D. ensifolium* showed an intact start codon for the *ndh*C gene but suffered a severe truncation with the loss of ca. 50% of homologous bases compared to the functional *ndh*C gene of *M. coccinea* (KP205432, Kim et al., 2015).

In *D. pandanum* 1, *D. stenocheilum* 1, and *D. variegatum* the *ndh*F gene was putatively lost (**Figure 6**). All other samples possessed severely truncated *ndh*F genes with absent start codons and multiple internal stop codons. The *ndh*H gene was present in *D. pandanum* 1 possessing a length of 1,176 bp (99.2% of homologous length compared to *M. coccinea* (KP205432, Kim et al., 2015). In most other samples, the *ndh*H gene was degraded possessing several stop codons. *D. stenocheilum* 1 and *D. variegatum* lost the *ndh*H gene (**Figure 5**, **Figure 6**).

Moreover, *ndh*E, *ndh*I and *ndh*A were found to be putatively lost in the plastome of *D. variegatum* and *D. pandanum* 1, respectively (**Figure 5**, **Figure 6**). No gene loss occurred for *ndh*B, *ndh*D and *ndh*J, however all three genes exhibited various degrees of degradation within all examined *Dipodium* plastomes and were either moderately or severely pseudogenised due to internal stop codons or frame-shift mutations.

The *ndh*D gene was found to have undergone the fewest degradation processes in regards of gene length which was largely conserved ranging from 1,122 bp (*D. campanulatum* 1) to 1,521 bp (*D. pulchellum*) and in most plastomes *ndh*D possessed the alternative start codon ACG (Threonine). Furthermore, almost all *Dipodium* plastomes showed the canonical AUG (Methionine) start codon for *ndh*A, *ndh*B, *ndh*E, and *ndh*I. The intron-containing *ndh*A and *ndh*B genes exhibited the strongest degradation (i.e., large deletions) within the intron regions and the downstream exon in all *Dipodium* samples. Exon1 of *ndh*B was almost complete and in-frame for most plastomes and showed only one point mutation (from A to C) which resulted in a stop codon at amino acid position 68 (after 201 bp from the beginning of the first exon in *ndh*B).

Within different *Dipodium* complexes the patterns for putative *ndh* gene losses and severe or moderate pseudogenisations were similar for examined plastomes of *D. hamiltonianum* and *D. interaneum.* Both plastomes putatively lost *ndh*G and *ndh*K and showed severe pseudogenisations of *ndh*A, *ndh*B, *ndh*C, *ndh*F, *ndh*H, *ndh*J and a moderately pseudogenised *ndh*E gene, but differed in level of putative pseudogenisation of *ndh*D and *ndh*I (**Figure 6**).

Other similarities were found in the *D. roseum* complex, in which the *ndh*D gene was moderately pseudogenised in all samples. Almost all samples of the *D. roseum* complex, except for *D. roseum* 2, harboured moderately pseudogenised *ndh*E and *ndh*I genes.

Within the same species, only *D.* aff. *roseum* 3 and *D.* aff. *roseum* 4 showed the same pattern of *ndh* gene loss and level of degradation which was also present in the plastome of *D. pardalinum* 2. Within the *D. stenocheilum* complex, *D. stenocheilum* 1 putatively lost *ndh*I and *ndh*F.

Across other samples, only two plastome pairs (*D. ensifolium* and *D.* aff. *roseum* 1; *D. basalticum,* and *D. atropurpureum* 2) shared the same pattern of *ndh* gene loss and degradation. In comparison to all other species of examined *Dipodium* plastomes, *D. variegatum* independently lost *ndh*E and *ndh*I and *D. pandanum* 1 lost the *ndh*A gene (**Figure 6**).

## 4 Discussion

This is the first molecular study to elucidate interspecific relationships and divergence times in *Dipodium* and to examine plastid genome degradation within a mycoheterotrophic orchid genus of the Australasian flora in a phylogenomic context.

### 4.1 Phylogenetic placement and infrageneric relationships of *Dipodium*

This phylogenomic study based on 68 plastid loci provided strong support for the monophyly of *Dipodium* and its phylogenetic placement as an early diverging lineage within tribe Cymbidieae. Previous phylogenetic studies included only one or two species of *Dipodium* which precluded assessment of the monophyly of the genus (Pridgeon et al., 2009: Chase et al., 2015; Górniak et al. (2010); Batista et al. (2014); Freudenstein and Chase (2015); Kim et al., 2020; Serna-Sánchez et al., 2021; McLay et al., 2023; Pérez-Escobar et al., 2023). Our study resolved *Dipodium* as the diverging early within Cymbidieae after subtribe Cymbidiinae with strong support and thus confirmed previous molecular phylogenetic studies in support of recognition of *Dipodium* at subtribal level as Dipodiinae (Li et al., 2016; Serna-Sánchez et al., 2021; Kim et al., 2020; Pérez-Escobar et al., 2023).

This phylogenomic study present the first molecular evidence in support of the infrageneric classification of *Dipodium* into sect. *Dipodium* and sect. *Leopardanthus* (O’Byrne, 2014; O’Byrne, 2017; Jones, 2021), lending support to the diagnostic value of vegetative traits (i.e., the presence or absence of adventitious roots) in infrageneric classification of *Dipodium*. Section *Leopardanthus* is characterised by leafy species which possess adventitious roots, such as *Dipodium pandanum.* In contrast, sect. *Dipodium* comprises species without adventitious roots and includes all leafless species, the leafy species *D. ensifolium,* and the morphologically similar *D. gracile* from Sulawesi, the latter being only known from the type (destroyed) (O’Byrne, 2017). Our phylogenomic study supported the placement of the *D. ensifolium* in sect. *Dipodium,* resolved as sister to all leafless species in the section. However, further molecular study is warranted to ascertain the monophyly of the two sections based on an expanded sampling of sect. *Leopardanthus*.

Our phylogenomic study is the first to shed light on evolutionary relationships within sect. *Dipodium,* which was found to comprise six main lineages. The phylogenomic framework now allows assessment of useful diagnostic morphological traits to characterise main lineages within the section. For example, the yellow stem and flower colour of species of the *D. hamiltonianum* complex easily distinguishes this clade from other mycoheterotrophic orchids within sect. *Dipodium* (**Figure 1**; Jones, 2021). Stems of remaining mycoheterotrophic species of sect. *Dipodium* are mostly greenish to dark reddish or purplish, whereas flowers vary in color from pale white, pinkish to purplish (**Figure 1**, Barrett et al., 2022; Jones, 2021). Also, sepal and petal characters were found to differ among clades: for example, species of clade A, comprising the *D. hamiltonianum* and *D. stenocheilum* complexes, possess sepals and lateral petals that are markedly narrower compared to species of clade B (comprising the *D. punctatum* and *D. roseum* complex) and *D. ensifolium*, the first diverging lineage within the sect. *Dipodium* (**Figure 1**; **Figure 3**) (Barrett et al., 2022; Jones, 2021).

Phylogenetic divergence within the two species complexes in clade B, *i.e.*, the *D. punctatum* and the *D. roseum* complexes, was shallow overall and thus interspecific relationships in these two groups remained largely unclear (**Figure 3**). Previous morphological studies highlighted difficulties in species delimitation within the *D. punctatum* complex, in particular between *D. pulchellum* and *D. punctatum* (Jones, 2021). While *D. pulchellum* is morphologically very similar to *D. punctatum*, the two species are differentiated by the intensity of their flower colours, which are richer in *D. pulchellum* and paler in *D. punctatum* (Jones and Clements, 1987). However, a morphological study by Jones (2021) revealed that the strong floral coloration of *D. pulchellum* flowers was likely due to differences in environmental factors (i.e., soil type and rainfall regime) of growing sites and thus Jones (2021) proposed to synonymise *D. pulchellum* with *D. punctatum*.

Similar challenges in taxonomic delimitation based on flower colours are also evident within the *D. roseum* complex. The distribution of the more widespread species *D. roseum* largely overlaps with the distributions of *D. atropurpureum* and *D. pardalinum* (ALA, 2023). Besides a very similar growing habit, the flowers of the three species are very similar in shape and vary only slightly in coloration: *D. roseum* has bright, rosy flowers with small darker spots, *D. atropurpureum* possesses dark pinkish-purple to dark reddish-purple flowers with spots and blotches, and the flowers of *D. pardalinum* are pale pink to white with large reddish spots and blotches (**Figure 1**) (Jones, 2021). Taken together, the overlapping distribution, similar appearance, and very shallow genetic divergence found in the present study among species in the *D. roseum* complex suggest that *D. atropurpureum* and *D. pardalinum* may be colour variations of *D. roseum*. Further molecular study with more highly resolving molecular techniques such as genotyping-by-sequencing is required to rigorously assess species delimitation within *Dipodium*.

### 4.2 Divergence-time estimations

Our divergence time estimations yielded results comparable to previous studies regarding the temporal diversification of major orchid clades (e.g., Givnish et al., 2015, Givnish et al., 2018; Kim et al., 2020; Serna-Sánchez et al., 2021; Zhang et al., 2023). Within Epidendroideae, this study confirmed that Cymbidieae was one of the most recently diverged tribes in Orchidaceae, consistent with previous studies (e.g., Givnish et al., 2015; Serna-Sánchez et al., 2021; Zhang et al., 2023). Stem and crown diversification of Cymbidieae were estimated to have commenced at ca. 42.2 Ma and 38.0 Ma respectively, which is similar to the estimates of Serna-Sánchez et al. (2021) and slightly younger than those of Zhang et al. (2023) (**Figure 4**, Supplementary Material 4 and 5).

Our study is the first to elucidate phylogenetic relationships and divergence times within *Dipodium*. Previously, only two studies included a representative of *Dipodium* (*D. roseum*, MN200368) in divergence-time estimations for Orchidaceae (Kim et al., 2020; Serna-Sánchez et al., 2021). These studies estimated the origin of *Dipodium* to ca. 17 Ma and ca. 31 Ma, respectively. Our study placed the divergence of *Dipodium* from the other subtribes in Cymbidieae to ca. 33.3 Ma in the early Oligocene which is closer to the findings of Serna-Sanchez et al. (2021). O’Byrne (2014) hypothesised that lineage divergence into sect. *Dipodium* and sect. *Leopardanthus* resulted from vicariance in conjunction with the break-up of Pangaea, in particular the separation of the Indian and Australian continental plates (O’Byrne, 2014). However, our divergence-time estimations show that *Dipodium* is far too young (< 33 Ma) to have been influenced by the break-up of Pangaea, which occurred from the early Jurassic and onwards. Lineage divergence of sect. *Dipodium* and sect. *Leopardanthus* were estimated to ca. 11.3 Ma in the late Miocene (**Figure 4**), when Australia had already assumed, approximately, its present geographical position. Rather, *Dipodium* is likely to have achieved its current distribution through range expansion between Australia and Southeast Asia across the Sunda-Sahul Convergence Zone (Joyce et al. 2021a), consistent with a general pattern of floristic exchange – the Sunda-Sahul Floristic Exchange - which was initiated as early as c. 30 Ma (Crayn et al., 2015; Joyce et al., 2021b). However, the data are insufficient at present to resolve the ancestral area of *Dipodium* and its main lineages. Further research is needed including an increased sampling to shed light on range evolution of *Dipodium* through ancestral range reconstruction.

Our results indicate that the Australian leafy species *D. ensifolium* diverged from the remainder of section *Dipodium* approximately 8.1 Ma (late Miocene) (**Figure 4**). The remainder of the sect. *Dipodium* clade, which includes all leafless, putatively fully mycoheterotrophic species, emerged ca. 7.3 Ma (late Miocene) followed by rapid diversification from ca. 4.3 Ma onwards (early Pliocene) (**Figure 4**). Thus, mycoheterotrophy has most likely evolved only once within *Dipodium*, on the Australian continent during the late Miocene-early Pliocene.

From the late Miocene-early Pliocene (ca. 5 Ma) climatic conditions in Australia became increasingly arid, leading to a decline of rainforest vegetation and expansion of open sclerophyllous forests (Quilty, 1994, Gallagher et al., 2003, Martin, 2006, He and Wang, 2021). By the end of the Pliocene Australia’s landscape was similar to the present day, with much of the continent a mosaic of open woody vegetation dominated by *Eucalyptus*, *Acacia* and Casuarinaceae (e.g., Martin 2006). The Pleistocene (ca. 2.58 – 0.012 Ma) was characterised by climatic oscillations which led to repeated forest expansion and contraction (Byrne, 2008). The evolution of mycoheterotrophy and the subsequent radiation of sect. *Dipodium* may have been facilitated by two factors: aridification in Australia favouring the reduction of leaf area to decrease water loss (O’Byrne, 2014), and the expansion of sclerophyll taxa and their mycorrhizal partners. Mycoheterotrophic *Dipodium* are assumed to share mycorrhizal fungi with Myrtaceae trees, especially *Eucalyptus*, (Bougoure and Dearnaley, 2005; Dearnaley and Le Brocque, 2006; Jones, 2021) which explosively diversified and came to dominate most Australian forests and presumably led to an increased diversity and abundance of suitable mycorrhizal partners for *Dipodium*. The rapid diversification of *Dipodium* from the Pleistocene onwards (ca. 3.2–0.3 Ma) (**Figure 4**) may have been driven by cycles of population fragmentation and coalescence in response to climatic oscillations.

### 4.3 Plastid genome evolution

#### 4.3.1 Plastome structural features and variations

In this study, whole plastome assemblies were generated for 24 *Dipodium* samples, including representatives of all leafless, putatively full mycoheterotrophs of sect*. Dipodium* found in Australia, one leafy photosynthetic species of sect. *Dipodium* (*D. ensifolium*) and one leafy photosynthetic species of sect. *Leopardanthus* (*D. pandanum*). The overall organisation and the plastid gene content is generally conserved in most examined *Dipodium* plastomes (**Figure 5**, **Table 2** and **3**). All examined plastomes showed the typical quadripartite structure of angiosperms (Ruhlman and Jansen 2014). However, some genomic features among several *Dipodium* plastomes were not conserved, including 1) differences in total genome length; 2) independent boundary shift IRB/SSC/IRA within the plastome of *D. pandanum* 1, *D. stenocheilum* 1, *D. variegatum*; 3) triplication of the *trn*V-GAC in the plastomes of *D. ammolithum, D. elegantulum*, *D. hamiltonianum*, *D. stenocheilum* 2, *D. interaneum* 4) the independent pseudogenisation of *cem*A in the plastome of *D.* aff. *roseum* 4 and *trn*D-GUC in the plastome of *D. campanulatum* 1; and 5) the pseudogenisation or loss to varying degrees of *ndh* genes (**Figure 5**, **Table 3**, Supplementary Material 7).

Total genome length of *Dipodium* plastomes displayed differences of around 10,000 bp between the smallest (142,949 bp; *Dipodium variegatum*) and largest plastomes (152,956 bp *Dipodium* aff. *roseum* 3) which correlated with level of *ndh* gene degradation. Some *Dipodium* plastomes were similar to the average size of orchid plastomes (152,442 bp) published on NCBI database (286 Orchidaceae chloroplast genome, accessed on June 13, 2022), however most plastomes were smaller (average size *Dipodium* plastomes: 148,703 bp; **Table 2**). Average GC contents in *Dipodium* was very similar to the average GC content of published orchid plastomes on NCBI database (ca. 36.8%; 286 Orchidaceae chloroplast genome, accessed on June 13, 2022) and all fell into the range of typical angiosperm plastomes (ca. 30–40%) (**Table 2**).

#### 4.3.2 Patterns of *ndh* gene degradation within *Dipodium*

In orchids, *ndh* gene losses and pseudogenisations which occurred in both autotrophic and heterotrophic species have been documented in various genera (e.g., Kim et al., 2015; Feng et al., 2016; Niu et al., 2017; Barrett et al., 2018, Barrett et al., 2019; Roma et al., 2018; Lallemand et al., 2019; Kim et al., 2020; Peng et al., 2022; Kim et al., 2023). This study is in line with these general findings in that *ndh* gene degradation was also observed within the orchid genus *Dipodium*. All chloroplast *ndh* genes in *Dipodium* plastomes exhibited varying degrees of putative pseudogenisation and loss, not a single *ndh* gene remained functional among the examined chloroplast genomes (**Table 3**, **Figure 5**, **Figure 6**). These findings include all plastomes of leafless putatively fully mycoheterotrophic species and of two autotrophic leafy species (*D. pandanum* and *D. ensifolium*) and thus suggest that all examined species, independently of their nutritional status, have lost the functionality of the plastid NADH dehydrogenase complex. Hence, the last common ancestor of extant *Dipodium* is likely to have lacked a functional NDH complex. Previous studies in Cymbidiinae, the first diverging lineage in Cymbideae, found that all species studied so far exhibited at least one degraded *ndh* gene (e.g., Yang et al., 2013; Kim and Chase 2017). As the next diverging lineage in Cymbidieae is *Dipodium*, this suggests that the degradation of *ndh* genes in Cymbidieae was likely a dynamic process from functional to non-functional. However, further research is needed e.g., ancestral state reconstructions of gene degradation with increased taxonomic sampling. The inclusion of more species among sect. *Leopardanthus* is warranted to clarify if some *ndh* genes have remained functional in some autotrophic species of sect. *Leopardanthus*.

Previous studies examined *ndh* gene loss at genus level and revealed an independent loss of function of the NADH dehydrogenase complex for several genera (e.g., Lin et al., 2015, Kim et al., 2015). However, comparative whole plastome studies examining gene degradation and loss among closely related mycoheterotrophic species are still scarce. For a better understanding of *ndh* gene degradation patterns this study investigated the degree of *ndh* gene degradation among closely related orchid species (**Figure 6**). Greatest degradation within *Dipodium* were found for *ndh*G which is putatively lost in almost all examined plastomes, except *D. stenocheilum* 1 which retained a putative severely pseudogenised *ndh*G (**Figure 6**). The *ndh*G gene is located within the SSC region. In general, it is well established that genes in the SSC region experience higher substitution rates compared to genes located within IR regions (Ruhlman and Jansen 2014). The latter is the case for *ndh*B which is located in the IRs and structurally more conserved in *Dipodium* compared to most *ndh* genes located in the SSC. The greatest degree of *ndh* gene degradation occurred in *D. variegatum* which putatively lost *ndh*C and *ndh*E–*ndh*I. All other plastomes putatively lost at least one to three *ndh* genes and showed different levels of degradation (**Figure 6**).

Interestingly, the level of *ndh* gene degradation varied even among closely related species within species complexes. For example, *D. stenocheilum* 1 independently lost *ndh*I and *ndh*F, whereas all other studied samples of the *D. stenocheilum* complex retained those two genes as moderately or severely pseudogenised (**Figure 6**). Different levels of gene degradation and loss were even found within the same species. For example, *D. atropurpureum* 1 lost *ndh*C whereas *D. atropurpureum* 2 retained a severely pseudogenised *ndh*C (**Figure 6**). Moreover, the study of Kim et al., (2020) included one individual of *D. roseum* which showed a different pattern of *ndh* gene loss and degradation to those found among the *D. roseum* samples of this study. *D. roseum* (MN200386) experienced complete loss of *ndh*A, *ndh*C–*ndh*I and *ndh*K, but retained pseudogenised *ndh*B and *ndh*J genes (Kim et al., 2020). These findings also agree with the recent comparative plastome study on *D. roseum* and *D. ensifolium*: *D. roseum* (OQ885084) has retained truncated *ndh*B, *ndh*D and *ndh*J genes, but completely lost *ndh*A, *ndh*C, *ndh*E– *ndh*I and *ndh*K (McLay et al., 2023).

Overall, some patterns of *ndh* gene degradation found in this study in *Dipodium* are similar, however many were unique for each individual examined. Hence, this suggests that sect. *Dipodium* has undergone a recent and active *ndh* gene degradation which strongly implies a relaxed evolutionary selective pressure for the retention of the NDH complex.

#### 4.3.3 IR/SSC junctions and IR instability

Orchidaceae plastomes frequently show an expansion/shift of the IR towards the SSC region (e.g., Kim et al., 2020). This instability of the IR/SSC junction is assumed to correlate with the deletion of *ndh*F and has resulted in a reduction of the SSC, as observed in several Orchidaceae plastomes (e.g., Kim et al., 2015; Niu et al., 2017; Dong et al., 2018; Roma et al., 2018) and in other land plant plastomes (e.g., Amaryllidaceae, Bignoniaceae, Orobanchaceae) (Thode and Lohmann 2019; Li et al., 2021; Könyves et al., 2021). This study revealed reduced SSC regions for most examined plastomes which correlated with the degradation of the *ndh* gene suite located in the SSC. Compared to typical SSC regions found in angiosperms (ca. 20 kb, Ruhlman and Jansen 2014), the smallest SSC region was reduced by ca. 7,900 bp (*D. variegatum*) and the largest SSC region was reduced by ca. 4,700 bp (*D. ensifolium*) (**Table 2**, **Figure 5**). However, a large expansion of the IR such as found in *Vanilla* and *Paphiopedilum* plastomes (Kim et al., 2015) was not found in *Dipodium* (IR sizes ranging between 24,436– 26,817 bp, **Table 2**).

In angiosperms, the *ycf*1 gene usually occupies ca. 1,000 bp in the IR (Sun et al., 2017, Kim et al., 2015). *Dipodium* plastomes in this study displayed varying positions of *ycf*1 within the IR. In plastomes in which the *ndh*F gene was completely lost or severely truncated, the portion of *ycf*1 within the IRA was mostly shorter compared to plastomes which contained moderately truncated *ndh*F genes (**Figure 5**). These results are similar with findings of Kim et al., (2015), a study which compared the locations of the IR/single-copy region junctions among 37 orchid plastomes and closely related taxa in Asparagales. In at least three plastomes (*D. pandanum* 1, *D. stenocheilum* 1, *D. variegatum*) *ndh*F was independently lost, the SSC/IRB junction was shifted into the spacer region near the *rpl*32 gene in direct adjacency to the partially duplicated *ycf*1 fragment (**Figure 5**, D, b–d). These findings suggest the deletion of *ndh*F correlated with the shift of the SSC/IRB junction. Interestingly, the boundaries between SSC and IR regions were found to be variable even among closely related species e.g., in *Cymbidium*. Some species in *Cymbidium* showed similar patterns of IR/SSC shifts (Kim and Chase 2017) as found in *Dipodium*.

In at least five plastomes (*D. ammolithum, D. elegantulum*, *D. hamiltonianum*, *D. interaneum, D. stenocheilum* 2) the *trn*V-GAC gene was triplicated (i.e., duplicated *trn*V-GAC version in close proximity to each other either in IRA or IRB) (**Figure 5**, B, C; **Table 3**). To the best of our knowledge, similar tRNA duplication patterns within the IR regions have not yet been found in any other Orchidaceae plastome. However, a recent study on plastomes of the angiosperm genus *Medicago* (Wu et al., 2021) yielded similar patterns. Wu et al. (2021) have found three copies of the *trn*V-GAC gene in the plastomes of two closely related species within the IR (*M. archiducis-nicolai* and *M. ruthenica*) which were linked to forward and tandem repeats. Interestingly, Wu et al. (2021) findings support the hypothesis that repetitive sequences lead to genomic rearrangements and thus affect plastome stability. This may also apply for some *Dipodium* plastomes. However, to rule out any technical issues throughout the NGS process and to validate findings of duplicated tRNAs (and above-mentioned boundaries of IR/SC regions), PCR amplification of affected regions should be carried out in future studies. However, in strong support of tRNA duplication is their independent presence within the IR of five plastomes among individuals of the same species complexes (*D. stenocheilum* complex and *D. hamiltonianum* complex). However, an increased sampling is necessary to better understand the impacts of genomic rearrangements due to repetitive sequences and thus plastome instability in *Dipodium*.

#### 4.3.4 Evolution of mycoheterotrophy and associated plastome degradation in *Dipodium*

Heterotrophic plants are remarkable survivors, exhibiting often curious morphological, physical, or genomic modifications. Multiple heterotrophs were found to have suffered plastid genome degradations due to relaxed pressure on photosynthetic function. In recent years, evidence has accumulated that plastid genomes have undergone gene degradation in the evolutionary transition from autotrophy to heterotrophy (e.g., Graham et al., 2017; Barrett et al., 2019; Wicke et al., 2016). Among these, the first stage is the loss and pseudogenisation of genes involved in encoding the NDH complex. Interestingly, all examined plastomes of *Dipodium* have lost or pseudogenised all 11 *ndh* genes regardless of their nutritional status (**Figure 6**). Two photosynthetic species with green leaves were included in this study, *D. pandanum* (sect. *Leopardanthus*) and *D. ensifolium* (sect. *Dipodium*). Degradation in *ndh* genes among photosynthetic species is not surprising and was frequently reported in previous plastome studies in land plants. The large-scale study on Orchidaceae plastomes of Kim et al., (2020) observed *ndh* gene pseudogenisation and losses among species in many epiphytes and several terrestrials which have retained their photosynthetic capacity. The NDH complex is thought to mediate the Photosystem I cyclic electron transport, fine-tunes photosynthetic processes and alleviates photooxidative stress (e.g., Yamori et al., 2015; Peltier et al., 2016; Sabater 2021). *D. pandanum* is a terrestrial or climbing epiphytic orchid and highly localised in rainforest habitats, whereas the terrestrial *D. ensifolium* grows in open forests and woodlands (Jones, 2021), thus both species seem to prefer shaded understory habitats. For epiphytic or terrestrial plants living in low-light habitats it has been proposed that the NDH complex may not be essential anymore (e.g., Barrett et al., 2019). One reason for this may be that they are less exposed to photooxidative stress (e.g., Feng et al., 2016; Barrett et al., 2019). However, the NDH complex is composed of 11 chloroplast encoded subunits and additional subunits encoded by the nucleus (e.g., Peltier et al., 2016). It has been established that genomic material was repeatedly exchanged between the nucleus, mitochondrion, and chloroplast in the evolutionary course of endosymbiosis. Thus, previous studies examined whether genes were transferred from the chloroplast to the nucleus and/or mitochondrion genome or whether nuclear genes for the NDH complex suffered under degradation. Indeed, Lin et al., (2015) reported *ndh* fragments within the mitochondrial genomes of orchids, however no copies were found in the nuclear orchid genomes. Similar findings were reported from the orchid genus *Cymbidium* (Kim and Chase et al., 2017). However, further studies are needed to determine whether *ndh* gene transfer into the nucleus or mitochondrion may play a role within *Dipodium*. The proposed subsequent next steps toward (myco-) heterotrophy is the functional loss of photosynthetic genes (e.g., *psa*, *psb*, *pet*, *rbc*L or *rpo*) followed by genes for the chloroplast ATP synthase and genes with other function such as housekeeping genes (e.g., *mat*K, *rpl*, *rnn* (e.g., Graham et al., 2017; Barrett et al., 2019). Most examined *Dipodium* plastomes displayed no additional plastid gene degradation besides *ndh* gene degradation, except in *D*. aff. *roseum* 4 where *cem*A was pseudogenised and in *D. campanulatum* 1 where the *trn*D-GUC gene was pseudogenised (**Table 3**). The *cem*A gene encodes the chloroplast envelope membrane protein and was found to be non-essential for photosynthesis, however *cem*A-lacking mutants of the green alga *Chlamydomonas* were found to have a severely affected carbon uptake (Rolland et al., 1997) and may therefore be classified as directly involved in photosynthesis. Transfer RNA genes (*trn*) are involved in the translation process and categorised as ‘housekeeping’ genes (e.g., Graham et al., 2017; Wicke and Naumann 2018; Barrett et al., 2019). Moreover, similar gene degradation patterns were found in the plastomes of *D. roseum* (MN200386, Kim et al., 2020 and OQ885084, Mclay et al., 2023) and *D. ensifolium* (OQ885084, Mclay et al., 2023), which functionally lost all *ndh* genes. However, most photosynthesis related genes in the plastomes of *Dipodium* were found to be functional. Thus, mycoheterotrophic species of *Dipodium* display evidence of being at the beginning of plastid gene degradation, in contrast with the majority of fully mycoheterotrophic orchids which are in more advanced stages of degradation, e.g. *Cyrtosia septentrionalis* (Kim et al., 2019), *Epipogium* (Schelkunov et al., 2015), and *Rhizanthella* (Delannoy et al., 2011). On the other hand, mycoheterotrophs such as *Corallorhiza trifida* (Barrett et al., 2018), *Cymbidium macrorhizon* (Kim et al., 2017), *Hexalectris grandiflora* (Barrett et al., 2019) and *Limodorum abortivum* (Lallemand et al., 2019) display functionally losses within the plastid *ndh* genes only and some species among them additionally lost one or two other genes, similar to findings in *Dipodium*. Interestingly, most of these species are leafless, but considered putatively partially mycoheterotrophic. Suetsugu et al. (2018) demonstrated that the leafless green orchid *Cymbidium macrorhizon* contains chlorophyll and can fix significant quantities of carbon during the fruit and seed production phase and thus, is photosynthetically active. Chlorophyll is present in *Corallorhiza trifida* also, but this green, leafless coralroot is an inefficient photosynthesiser (Barrett et al., 2014). Some species among leafless orchids within sect. *Dipodium* (e.g., *D. elegantulum*, *D. stenocheilum*, *D. variegatum*) appear green on stems (**Figure 1**, Jones 2021), which suggests they may contain some chlorophyll and be able to photosynthesise. Coupled with relatively mild plastid gene degradation compared to other fully mycohetrotrophic orchids, this suggests some leafless species among sect. *Dipodium* may be partially mycoheterotrophic rather than fully mycoheterotrophic as has been hypothesised for *D. roseum* (Kim et al. 2020; McLay et al. 2023). However, no studies so far have examined whether leafless species among sect. *Dipodium* contain chlorophyll and whether they are capable to carry out photosynthesis at sufficient rates. Therefore, more research is needed to assess the trophic status, including analysis of chlorophyll quantities and the ratio of photosynthetic carbon to fungal carbon for *Dipodium*.

Compared with recently published studies on mycoheterotrophic orchids such as *Corallorhiza* and *Hexalectris* (Barret et al. 2018; Barret et al. 2019) which incorporated divergence time estimations, plastomes of *Dipodium* showed the least degradation. *Hexalectris* crown age was estimated to ca. 24 Ma and plastomes of mycoheterotophs were more degraded compared to mycoheterotrophic plastomes of *Corallorhiza* which diversified ca. 9 Ma onwards (Barret et al. 2018; Barret et al. 2019). *Dipodium* diversified in the late Miocene ca. 11 Ma, and the mycoheterotrophic lineage divergent from the autotrophic lineage ca. 8.1 Ma which is slightly younger compared to *Corallorhiza*. Hence, time of divergence may play a role in the degree of degradation of *Dipodium* plastomes which show an early stage of plastome degradation compared to older diverging mycoheterotrophic lineages that are in more advanced stages of plastome degradation.

## 5 Conclusion

This molecular phylogenomic comparative study clarified evolutionary relationships and divergence times of the genus *Dipodium* and provided support for two main lineages within *Dipodium,* corresponding to the morphologically defined sect. *Dipodium* and sect. *Leopardanthus*. Phylogenetic analysis resolved the leafy autotroph *D. ensifolium* as being part of sect. *Dipodium* and found to be in sister group position to all leafless species in sect. *Dipodium*. Divergence-time estimations placed the divergence of the leafy species *D. ensifolium* from the remainder of section *Dipodium* in the late Miocene. Shortly after, the remaining clade including all leafless, putatively full mycoheterotrophic species within sect. *Dipodium* emerged ca. 7.3 Ma in the late Miocene followed by rapid species diversification from ca. 4.3 Ma onwards in the early Pliocene. Thus, this study indicates that mycoheterotrophy has most likely evolved only once on the Australian continent within *Dipodium* during the late Miocene, and that the ancestors of putatively full mycoheterotrophic species may have had green leaves. Among the examined plastomes, all plastid *ndh* genes were pseudogenised or physically lost, regardless of the individual’s nutrition strategy (i.e., autotroph versus mycoheterotroph). Thus, this study provides molecular evidence of relaxed evolutionary selective pressure on the retention of the NADH dehydrogenase complex. Mycoheterotrophic species among sect. *Dipodium* retained a full set of other functional photosynthesis-related genes and exhibited an early stage of plastid genome degradation. Hence, leafless species of sect. *Dipodium* may potentially be rather partially mycoheterotrophic than fully mycoheterotrophic.

To further disentangle evolutionary relationships in *Dipodium,* future studies based on nuclear data such as derived from target capture sequencing and with a denser sampling at population level are warranted. Moreover, the inclusion of a denser sampling of sect. *Leopardanthus* is warranted to clarify if some *ndh* genes may have remained functional in some of the autotrophic species of sect. *Leopardanthus.* To obtain further insights into the nutritional strategies in *Dipodium*, future studies should assess the trophic status of mycoheterotrophic species in *Dipodium* based on physiological data such as from the analysis of chlorophyll quantities and the ratio of photosynthetic carbon to fungal carbon for *Dipodium*. The Australian orchid flora harbours many more remarkable mycoheterotrophic lineages (e.g., *Danhatchia*) which offer the opportunity to further explore the evolutionary pathways to mycoheterotrophy and associated plastid genome evolution. The inclusion of autotrophic plants into comprehensive plastid phylogenetic analyses could broaden the understanding of the significance of observed *ndh* gene degradation patterns within Orchidaceae.

## Author Contributions

**Conceptualisation:** SG, KN, MAC. **Methodology:** SG, KN, SJB; **Data curation:** SG, KN, MAC. **Formal analysis:** SG, SJB. **Funding acquisition:** KN, DMC, MAC, SG. **Investigation:** SG, KN, MAC, SJB, JAN, VSP, PMS. **Visualisation:** SG. **Writing – original draft:** SG. **Writing – review & editing**: SG, KN, MAC, SJB, JAN, VSP, PMS, DMC.

## Acknowledgements

The authors acknowledge the contribution of Bioplatforms Australia (enabled by NCRIS) in the generation of data used in this publication. We acknowledge K. Alcock, M.D. Barrett, C. Bower, C.P. Brock, R. Crane, D.M. Crayn, W. Dowling, J. Egan, B. Gray, C. Houston, M. Jacobs, D.L. Jones, P.D. Jones, C.D. Kilgour, L. Lawler, K.R. McDonald, I. Morris, D.E. Murfet, J. Taylor for collection of plant material used in this study.

## Supplementary Material

**Supplementary Material 1.** Details of samples included in phylogenetic analysis and divergence-time estimations.

**Supplementary Material 2. a.** Details of plastid loci included in alignment of ML-phylogenetic and divergence-time estimations. **b**. Parsimony informative sites (Pi) for each plastid gene.

**Supplementary Material 3.** ML-Phylogenetic tree of Orchidaceae.

**Supplementary Material 4. a.** Model comparison by AICM (Akaike Information Criterion by MCMC) **b.** Comparison divergence-time estimations of major Orchidaceae linages (subfamilies), the tribe Cymbidieae and subtribe Dipodiinae.

**Supplementary Material 5**. Maximum-clade-credibility tree from Bayesian divergence-time estimations of Orchidaceae.

**Supplementary Material 6.** Summary of assembly features of 24 newly generated *Dipodium* plastomes.

**Supplementary Material 7.** Circular plastome maps of 24 newly generated *Dipodium* plastomes.

## References

ALA (2023). Atlas of Living Australia. Available at: https://www.ala.org.au (Accessed August,8 2023).

APC (2023): Australian Plant Census. Available at: biodiversity.org.au/nsl/services/search/taxonomy (Accessed May 5, 2023).

Altschul, S.F., Gish, W., Miller, W., Myers, E.W., Lipman, D.J. (1990). Basic local alignment search tool. J. Mol. Biol. 215, 403–410. 10.1016/S0022-2836(05)80360-2

Bankevich, A., Nurk, S., Antipov, D., Gurevich, A.A., Dvorkin, M., Kulikov, A.S., Lesin, V.M., Nikolenko, S.I., Pham, S., Prjibelski, A.D., Pyshkin, A.V., Sirotkin, A.V., Vyahhi, N., Tesler, G., Alekseyev, M.A., Pevzner, P.A. (2012). SPAdes: A new genome assembly algorithm and its applications to single-cell sequencing. J. Comput. Biol. 19, 455–477. 10.1089/cmb.2012.0021

Barrett, C.F., Freudenstein, J.V., Li, J., Mayfield-Jones, D.R., Perez, L., Pires, J.C., Santos, C. (2014). Investigating the path of plastid genome degradation in an early-transitional clade of heterotrophic orchids, and implications for heterotrophic angiosperms. Mol. Biol. Evol. 31, 3095–3112. 10.1093/molbev/msu252

Barrett, C.F., Wicke, S., Sass, C. (2018). Dense infraspecific sampling reveals rapid and independent trajectories of plastome degradation in a heterotrophic orchid complex. New Phytol. 218, 1192–1204. 10.1111/nph.15072

Barrett, C.F., Sinn, B.T., Kennedy, A.H. (2019). Unprecedented parallel photosynthetic losses in a heterotrophic orchid genus. Mol. Biol. Evol. 36, 1884–1901. 10.1093/molbev/msz111

Barrett, R. L., Barrett, M.D., Clements, M.A. (2022). A revision of Orchidaceae from the Kimberley region of Western Australia with new species of tropical *Calochilus* and *Dipodium*. Telopea 25, 203–270. 10.7751/telopea15711

Batista, J.A.N., Mota, A.C.M., Proite, K., Bianchetti, L.D.B., Romero-González, G.A., Huerta, H., Salazar, G.A. (2014). Molecular phylogenetics of neotropical *Cyanaeorchis* (Cymbidieae, Epidendroideae, Orchidaceae): geographical rather than morphological similarities plus a new species. Phytotaxa 156, 251–272. 10.11646/phytotaxa.156.5.1

Bolger, A.M., Lohse, M., Usadel, B. (2014). Trimmomatic: a flexible trimmer for Illumina sequence data. Bioinformatics 30, 2114–2120. 10.1093/bioinformatics/btu170

Bouckaert, R., Heled, J., Suchard, M.A., Rambaut, A., Drummond, A.J. (2014). BEAST 2: A software platform for Bayesian evolutionary analysis. PLOS Comput. Biol. 10, 4. 10.1371/journal.pcbi.1003537

Bouckaert, R., Vaughan, T.G., Barido-Sottani, J., Duchêne, S., Fourment, M., Gavryushkina, A., Heled, J., Jones, G., Kühnert, D., De Maio, N., Matschiner, M., Mendes, F.K., Müller, N.F., Ogilvie, H.A., du Plessis, L., Popinga, A., Rambaut, A., Rasmussen, D., Siveroni, I., Suchard, M.A., Wu, C.H., Xie, D., Zhang, C., Stadler, T., Drummond, A.J. (2019). BEAST 2.5: An advanced software platform for Bayesian evolutionary analysis. PLOS Comput. Biol. 15, 4. 10.1371/journal.pcbi.1006650

Bougoure, J.J., Dearnaley, J.D.W (2005). The fungal endophytes of *Dipodium variegatum* (Orchidaceae). Australasian Mycologist 24, 15–19.

Bushnell, B. (2014). BBMap: A fast, accurate, splice-aware aligner. Lawrence Berkeley National Laboratory, Berkeley, CA (United States).

Braukmann, T.W.A., Broe, M.B., Stefanović, S. and Freudenstein, J.V. (2017). On the brink: the highly reduced plastomes of nonphotosynthetic Ericaceae. New Phytol. 216, 254–266. 10.1111/nph.14681

Byrne, M., Yeates, D.K., Joseph, L., Kearney, M., Bowler, J., Williams, M.A.J., Cooper, S., Donnellan, S.C., Keogh, J.S., Leys, R., Melville, J., Murphy, D.J., Porch, N. and Wyrwoll, K.-H. (2008). Birth of a biome: insights into the assembly and maintenance of the Australian arid zone biota. Mol. Ecol. 17, 4398–4417. 10.1111/j.1365-294X.2008.03899.x

Chase, M.W., Cameron, K.M., Freudenstein, J.V., Pridgeon, A.M., Salazar, G., Van den Berg C., Schuitemanp, A. (2015). An updated classification of Orchidaceae. Bot. J. Linn. Soc. 177, 151–174. 10.1111/boj.12234

Christenhusz, M.J.M, Byng, J.W. (2016). The number of known plants species in the world and its annual increase. Phytotaxa 261, 201. 10.11646/phytotaxa.261.3.1.

Crayn, D.M., Costion, C., Harrington, M.G. (2015). The Sahul–Sunda floristic exchange: dated molecular phylogenies document Cenozoic intercontinental dispersal dynamics. J. Biogeog. 42, 11–24. 10.1111/jbi.12405

Dearnaley, J.D.W., Le Brocque, A.F. (2006). Molecular identification of the primary root fungal endophytes of *Dipodium hamiltonianum* (Orchidaceae). Aust. J. Bot. 54, 487. 10.1071/BT05149

Delannoy, E., Fujii, S., Colas des Francs-Small, C., Brundrett, M., Small, I. (2011). Rampant gene loss in the underground orchid *Rhizanthella gardneri* highlights evolutionary constraints on plastid genomes. Mol. Biol. Evol. 28, 2077–2086. 10.1093/molbev/msr028

Douglas, J., Zhang, R., Bouckaert, R. (2021). Adaptive dating and fast proposals: Revisiting the phylogenetic relaxed clock model. PLoS Comput. Biol. 17 (2). 10.1371/journal.pcbi.1008322

Dong, W.L., Wang, R.N., Zhang, N.Y., Fan, W.B., Fang, M.F., Li, Z.H. (2018). Molecular evolution of chloroplast genomes of orchid species: insights into phylogenetic relationship and adaptive evolution. Int. J. Mol. Sci. 19, 716. 10.3390/ijms19030716

Drummond, A.J., Rambaut, A. (2007). BEAST: Bayesian evolutionary analysis by sampling trees. BMC Evol. Biol. 7, 214. 10.1186/1471-2148-7-214

Fabozzi, F.J., Focardi, S.M., Rachev, S.T., Arshanapalli, B.G. (2014). Appendix E. Model Selection Criterion: AIC and BIC in “The basics of financial econometrics: tools, concepts, and asset management applications.” John Wiley & Sons, Inc. 399–403.

Feng, Y.L., Wicke, S., Li, J.W., Han, Y., Lin, C.S., Li, D.Z., Zhou, T.T., Huang, W.C., Huang, L.Q., Jin, X.H. (2016). Lineage-specific reductions of plastid genomes in an orchid tribe with partially and fully mycoheterotrophic species. Genome Biol. Evol. 8, 2164–2175. 10.1093/gbe/evw144

Freudenstein, J.V., Chase, M.W. (2015). Phylogenetic relationships in Epidendroideae (Orchidaceae), one of the great flowering plant radiations: progressive specialization and diversification. Ann. Bot. 115, 665–681. 10.1093/aob/mcu253

Gallagher, S.J., Greenwood, D.R., Taylor, D., Smith, A.J., Wallace, M.W., Holdgate, G.R. (2003). The Pliocene climatic and environmental evolution of southeastern Australia: Evidence from the marine and terrestrial realm. *Palaeogeogr., Palaeoclimatol.*, Palaeoecol. 193, 349–382. 10.1016/S0031-0182(03)00231-1

Gernhard, T. (2008). The conditioned reconstructed process. J. Theor. Biol. 253, 769–778. 10.1016/j.jtbi.2008.04.005

Givnish, T.J., Zuluaga, A., Spalink, D., Soto Gomez, M., Lam, V.K.Y, Saarela, J.M., Sass, C., Iles, W.J.D, De Sousa, D.J.L, Leebens-Mack, J., Chris Pires, J., Zomlefer, W.B., Gandolfo, M.A., Davis, J.I., Stevenson, D.W., De Pamphilis, C., Specht, C.D., Graham, S.W., Barrett, C.F., Ané, C. (2018). Monocot plastid phylogenomics, timeline, net rates of species diversification, the power of multi-gene analyses, and a functional model for the origin of monocots. Am. J. Bot. 105, 1888–1910. 10.1002/ajb2.1178

Givnish, T.J., Spalink, D., Ames, M., Lyon, S.P., Hunter, S.J., Zuluaga, A., Iles, W.J.D., Clements, M.A., Arroyo, M.T.K, Leebens-Mack, J., Endara, L., Kriebel, R., Neubig, K.M., Whitten, W.M., Williams, N.H., Cameron, K.M. (2015). Orchid phylogenomics and multiple drivers of their extraordinary diversification. Proc. Royal Soc. B 282, 20151553. 10.1098/rspb.2015.1553

Górniak, M., Paun, O., Chase, M.W. (2010). Phylogenetic relationships within Orchidaceae based on a low-copy nuclear coding gene, *Xdh*: Congruence with organellar and nuclear ribosomal DNA results. Mol. Phyl. Evol. 56, 784–795. 10.1016/j.ympev.2010.03.003

Graham, S.W., Lam, V.K.Y., Merckx, V.S.F.T (2017). Plastomes on the edge: the evolutionary breakdown of mycoheterotroph plastid genomes. New Phytol. 214, 48–55. 10.1111/nph.14398

Greiner, S., Lehwark, P., Bock, R. (2019). OrganellarGenomeDRAW (OGDRAW) version 1.3.1: expanded toolkit for the graphical visualization of organellar genomes. Nucleic Acids Res. 47, W59–W64. 10.1093/nar/gkz238

Guindon, S., Dufayard, J.-F., Lefort, V., Anisimova, M., Hordijk, W., Gascuel, O. (2010). New algorithms and methods to estimate maximum-likelihood phylogenies: Assessing the performance of PhyML 3.0. Syst. Biol. 59, 307–321. 10.1093/sysbio/syq010

He, Y., Wang, H. (2021). Terrestrial material input to the northwest shelf of Australia through the Pliocene-Pleistocene period and its implications on continental climates. Geophys. Res. Lett., 48, e2021GL092745. 10.1029/2021GL092745

Hoang, D.T., Chernomor, O., Von Haeseler, A., Minh, B.Q., Vinh, L.S. (2018). UFBoot2: Improving the Ultrafast Bootstrap approximation. Mol. Biol. Evol. 35, 518–522. 10.1093/molbev/msx281

Jacquemyn, H., Merckx, V.S.F.T (2019). Mycorrhizal symbioses and the evolution of trophic modes in plants (R Shefferson, Ed.). J. Ecol. 107, 1567–1581. 10.1111/1365-2745.13165

Jones, D.L. (2021). A complete guide to native orchids of Australia. Sydney: Reed New Holland Publishers.

Jones, D.L. and Clements, M.A. (1987). New orchid taxa from south-eastern Queensland. Proc. R. Soc. Queensland 98, 128.

Joyce, E.M., Crayn, D.M., Lam, V.K.Y, Gerelle, W.K., Graham, S.W., Nauheimer, L. (2018). Evolution of *Geosiris* (Iridaceae): historical biogeography and plastid-genome evolution in a genus of non-photosynthetic tropical rainforest herbs disjunct across the Indian Ocean. Aust. Syst. Bot. 31, 504–522. 10.1071/SB18028

Joyce, E.M, Thiele, K.R., Slik, J.W.F., Crayn, D.M. (2021a). Plants will cross the lines: climate and available land mass are the major determinants of phytogeographical patterns in the Sunda–Sahul Convergence Zone. Biol. J. Linn. Soc. 132, 374–387. 10.1093/biolinnean/blaa194

Joyce, E.M., Pannell, C.M, Rossetto, M., Yap, J.Y.S., Thiele, K.R., Wilson, P.D., Crayn, D.M. (2021b). Molecular phylogeography reveals two geographically and temporally separated floristic exchange tracks between Southeast Asia and Northern Australia. J. Biogeog. 48, 1213–1227. 10.1111/jbi.14072

Kalyaanamoorthy, S., Minh, B.Q., Wong, T.K.F, Von Haeseler, A., Jermiin, L.S. (2017). ModelFinder: fast model selection for accurate phylogenetic estimates. Nat. Methods 14, 587–589. 10.1038/nmeth.4285

Katoh, K., Misawa, K., Kuma, K., Miyata, T., Mafft (2002). A novel method for rapid multiple sequence alignment based on fast Fourier transform, Nucleic Acids Res. 30 (14), 3059–3066, 10.1093/nar/gkf436

Katoh, K., Standley, D.M. (2013). MAFFT Multiple Sequence Alignment Software Version 7: Improvements in performance and usability. Mol. Biol. Evol. 30, 772–780. 10.1093/molbev/mst010

Kim, H.T., Kim, J.S., Moore, M.J., Neubig, K.M., Williams, N.H., Whitten, W.M., Kim, J.H. (2015). Seven new complete plastome sequences reveal rampant independent loss of the *ndh* gene family across orchids and associated instability of the inverted repeat/small single-copy region Boundaries (S. Aceto, Ed.). PLOS ONE 10, e0142215. 10.1371/journal.pone.0142215

Kim, H.T., Chase, M.W. (2017). Independent degradation in genes of the plastid *ndh* gene family in species of the orchid genus *Cymbidium* (Orchidaceae; Epidendroideae). PLOS ONE 12, e0187318. 10.1371/journal.pone.0187318

Kim, Y.K., Kwak, M.H., Chung, M.G., Kim, H.W., Jo, S., Sohn, J.Y., Cheon, S.H., Kim, K.J. (2017) The complete plastome sequence of the endangered orchid *Cymbidium macrorhizon* (Orchidaceae). Mitochondrial DNA Part B 2, 725–727. 10.1080/23802359.2017.1390411

Kim, Y.K., Jo, S., Cheon, S.H., Joo, M.J., Hong, J.R., Kwak, M.H., Kim, K.J. (2019). Extensive losses of photosynthesis genes in the plastome of a mycoheterotrophic orchid, *Cyrtosia septentrionalis* (Vanilloideae: Orchidaceae). Genome Biol. Evol. 11(2), 565–571. 10.1093/gbe/evz024

Kim, Y.K., Jo, S., Cheon, S.H., Joo, M.J., Hong, J.R., Kwak, M., Kim, K.J. (2020). Plastome evolution and phylogeny of Orchidaceae, with 24 new sequences. Front. Plant Sci. 11, 22. 10.3389/fpls.2020.00022

Kim, Y.K., Cheon, S.-H., Hong, J.-R., Kim, K.J. (2023). Evolutionary Patterns of the Chloroplast Genome in Vanilloid Orchids (Vanilloideae, Orchidaceae). Int. J. Mol. Sci. 24, 3808. 10.3390/ijms24043808

Klimpert, N.J., Mayer, J.L.S., Sarzi, D.S., Prosdocimi, F., Pinheiro, F., Graham, S.W. (2022). Phylogenomics and plastome evolution of a Brazilian mycoheterotrophic orchid, *Pogoniopsis schenckii*. Am. J. Bot. 109(12), 2030–2050. 10.1002/ajb2.16084

Könyves, K., Bilsborrow, J., Christodoulou, M.D., Culham, A., David, J. (2021). Comparative plastomics of Amaryllidaceae: inverted repeat expansion and the degradation of the *ndh* genes in *Strumaria truncate* Jacq. PeerJ 9, e12400. 10.7717/peerj.12400

Lam, V.K.Y., Gomez, M.S., Graham, S.W. (2015). The highly reduced plastome of mycoheterotrophic *Sciaphila* (Triuridaceae) is colinear with its green relatives and is under strong purifying selection, Genome Biol. Evol. 7, 2220–2236. 10.1093/gbe/evv134

Lallemand, F., Logacheva, M., Le Clainche, I., Bérard, A., Zheleznaia, E., May, M., Jakalski, M., Delannoy, É., Le Paslier, M.C., Selosse, M.A. (2019). Thirteen new plastid genomes from mixotrophic and autotrophic species provide insights into heterotrophy evolution in Neottieae orchids. Genome Biol. Evol. 11, 2457–2467. 10.1093/gbe/evz170

Li, M.H., Zhang, G.Q., Liu, Z.J., Lan, S.R. (2016). Subtribal relationships in Cymbidieae (Epidendroideae, Orchidaceae) reveal a new subtribe, Dipodiinae, based on plastid and nuclear coding DNA. Phytotaxa 246, 37. 10.11646/phytotaxa.246.1.3

Li, X., Yang, J.B., Wang, H., Song, Y., Corlett, R.T., Yao, X., Li, D.Z., Yu, W.B. (2021). Plastid NDH pseudogenization and gene loss in a recently derived lineage from the largest hemiparasitic plant genus *Pedicularis* (Orobanchaceae). Plant Cell Physiol. 62, 971–984. 10.1093/pcp/pcab074

Li, Z.H., Jiang, Y., Ma, X., Li, J.W., Yang, J.B., Wu, J.Y., Jin, X.H. (2020). Plastid genome evolution in the subtribe Calypsoinae (Epidendroideae, Orchidaceae) (J Archibald, Ed.). Genome Biol. Evol. 12, 867–870. 10.1093/gbe/evaa091

Lin, C.S., Chen, J.J., Huang, Y.T., Chan, M.T., Daniell, H., Chang, W.J., Hsu, C.T., Liao, D.C., Wu, F.H., Lin, S.Y., Liao, C.F., Deyholos, M.K., Wong, G.K., Albert, V.A., Chou, M.L., Chen, C.Y., Shih, M.C. (2015). The location and translocation of *ndh* genes of chloroplast origin in the Orchidaceae family. Sci. Rep. 12; 5:9040. 10.1038/srep09040

Logacheva, M.D., Schelkunov, M.I., Aleksey, A. P. (2011). Sequencing and analysis of plastid genome in mycoheterotrophic orchid *Neottia nidus*-*avis*. Genome Biol. Evol. 3, 1296–1303. 10.1093/gbe/evr102

Martin, H.A. (2006). Cenozoic climatic change and the development of the arid vegetation in Australia. J. Arid Environ. 66, 533–563. 10.1016/j.jaridenv.2006.01.009

McLay, T.G. B., Bayly, M. J., Whitehead, M. R., Fowler, R. M. (2023). Retention of an apparently functional plastome in an apparently mycoheterotrophic orchid, *Dipodium roseum* D.L.Jones & M.A.Clem. (Orchidaceae). Aust. J. Bot. 71, 306–317. 10.1071/BT22075

Merckx, V.S.F.T. (2013). *Mycoheterotrophy: an introduction* in *Mycoheterotrophy. The Biology of Plants Living on Fungi*. Berlin: Springer-Verlag 297–342. 10.1007/978-1-4614-5209-6

Minh, B.Q., Schmidt, H.A., Chernomor, O., Schrempf, D., Woodhams, M.D., Von Haeseler, A., Lanfear, R. (2020). IQ-TREE 2: New Models and efficient methods for phylogenetic inference in the genomic era. Mol. Biol. Evol. 37, 1530–1534. 10.1093/molbev/msaa015

Nargar, K., O’Hara, K., Mertin, A., Bent, S.J., Nauheimer, L., Simpson, L., Zimmer, H., Molloy, B.P.J., Clements, M.A. (2022). Evolutionary relationships and range evolution of greenhood orchids (subtribe Pterostylidinae): Insights from plastid phylogenomics. Front. Plant Sci. 13:912089. 10.3389/fpls.2022.912089

NCBI (2022), National Library of Medicine. Available at: https://www.ncbi.nlm.nih.gov (Accessed June 2, 2022)

Nguyen, L.T., Schmidt, H.A., Von Haeseler, A., Minh, B.Q. (2015). IQ-TREE: A fast and effective stochastic algorithm for estimating maximum-likelihood phylogenies. Mol. Biol. Evol. 32, 268–274. 10.1093/molbev/msu300

Niu, Z., Xue, Q., Zhu, S., Sun, J., Liu, W., Ding, X. (2017). The complete plastome sequences of four orchid species: Insights into the evolution of the Orchidaceae and the utility of plastomic mutational hotspots. Front. Plant Sci. 8, 715. 10.3389/fpls.2017.00715

O’Byrne, P. (2014). On the evolution of *Dipodium* R. Br. Reinwardtia 14 (1), 123–132 https://e-journal.biologi.lipi.go.id/index.php/reinwardtia/article/view/402 (Accessed November 22, 2021)

O’Byrne, P. (2017). A taxonomic revision of *Dipodium* section *Leopardanthus*. Malesian Orchid Journal 19, 5–142.

Peltier, G., Aro, E-M., Shikanai, T. (2016). NDH-1 and NDH-2 plastoquinone reductases in oxygenic photosynthesis. Ann. Rev. Plant Biology 67, 55–80. 10.1146/annurev-arplant-043014-114752

Peng, H.-W., Lian, L. Zhang, J., Erst A.S., Wang, W. (2022). Phylogenomics, plastome degradation and mycoheterotrophy evolution of Neottieae (Orchidaceae), with emphasis on the systematic position and Loess Plateau-Changbai Mountains disjunction of *Diplandrorchis*. BMC Plant Biol. 22, 5077. 10.1186/s12870-022-03906-0

Pérez-Escobar, O. A, Bogarín, D., Przelomska, N.A.S., Ackerman, J.D., Balbuena, J.A., Bellot, S. Bühlmann, R. P., Cabrera, B., Aguilar Cano, J., Charitonidou, M., Chomicki, G., Clements, M.A., Cribb, P., Fernández, M., Flanagan, N.S., Gravendeel, B., Hágsater, E., Halley, J.M., Hu, A.-Q., Jaramillo, C., Mauad, A.V., Maurin, O., Müntz, R., Leitch, I.J., Li, L., Negrao, R., Oses, L., Phillips, C., Rincon, M. Salazar-Chavez, G., Simpson, L., Smidt, E., Solano-Gomez, R., Parra-Sánchez, E., Tremblay, R.L., van den Berg, C., Villanueva, B.S., Zuluaga, A., Chase, M.W., Fay, M.F., Condamine, M.F., Forest, F. Nargar, N., Renner, S. S., Baker, W.J., Antonelli, A. (2023). The origin and speciation of orchids [*Preprint*]. Available at https://www.biorxiv.org/content/10.1101/2023.09.10.556973v3.full, (Accessed January 3, 2024). 10.1101/2023.09.10.556973

POWO (2023). Plants of the World Online. Facilitated by the Royal Botanic Gardens, Kew. Published on the Internet; http://www.plantsoftheworldonline.org/ (Accessed September 8, 2023).

Pridgeon, A.M., Cribb, P.J., Chase, M.W., Rasmussen, F.N. et al. (2009). Genera Orchidacearum, Volume 5, Epidendroideae (Part 2). Oxford University, Oxford (New York), 585.

Qu, X.J., Fan, S.J., Wicke, S., Yi, T.S. (2019). Plastome reduction in the only parasitic gymnosperm *Parasitaxus* is due to losses of photosynthesis but not housekeeping genes and apparently involves the secondary gain of a Large Inverted Repeat. Genome Biol. Evol. 11(10). 2789–2796. 10.1093/gbe/evz187

Quilty, P.G. (1994). The background: 144 million years of Australian paleoclimate and palaeogeography. In ‘History of the Australian vegetation: Cretaceous to recent’. 14–39. 2017 Edition. (Cambridge University Press: Cambridge, UK). 10.20851/australian-vegetation

Rambaut, A., Drummond, A.J., Xie, D., Baele, G., Suchard, M.A. (2018). Posterior summarization in Bayesian phylogenetics using Tracer 1.7. Syst. Biol. 67, 901–904. 10.1093/sysbio/syy032

Rolland, N., Dorne, A.J., Amoroso, G., Sültemeyer, D.F., Joyard, J., Rochaix, J.D. (1997). Disruption of the plastid *ycf*10 open reading frame affects uptake of inorganic carbon in the chloroplast of *Chlamydomonas*. EMBO J. 16, 6713–6726. 10.1093/emboj/16.22.6713

Roma, L., Cozzolino, S., Schlüter, P.M., Scopece, G., Cafasso, D. (2018). The complete plastid genomes of *Ophrys iricolor* and *O. sphegodes* (Orchidaceae) and comparative analyses with other orchids. PLOS ONE 13, e0204174. 10.1371/journal.pone.0204174

Ruhlman, T.A., Jansen, R.K. (2014). The plastid genomes of flowering plants. ‘Chloroplast Biotechnology’. Methods Mol. Biol. 3–38. 10.1007/978-1-62703-995-6_1

Sabater, B. (2021). On the edge of dispensability, the chloroplast *ndh* genes. Int. J. Mol. Sci. 22, 12505. 10.3390/ijms222212505

Schelkunov, M.I., Shtratnikova, V.Y., Nuraliev, M.S., Selosse, M.A., Penin, A.A., Logacheva, M.D. (2015). Exploring the limits for reduction of plastid genomes: A case study of the mycoheterotrophic orchids *Epipogium aphyllum* and *Epipogium roseum*. Genome Biol. Evol. 7, 1179–1191. 10.1093/gbe/evv019

Schlechter (1911). Lii. D. *gracile* Schltr. nov. spec. in Repertorium Specierum Novarum Regni Vegetabilis 10: 191. Berlin, Selbstverlag des Herausgebers 1912[1911].

Serna-Sánchez, M.A., Pérez-Escobar, O.A., Bogarín, D., Torres-Jimenez, M.F., Alvarez-Yela, A.C., Arcila-Galvis, J.E., Hall, C.F., De Barros, F., Pinheiro, F., Dodsworth, S., Chase, M.W., Antonelli, A., Arias, T. (2021). Plastid phylogenomics resolves ambiguous relationships within the orchid family and provides a solid timeframe for biogeography and macroevolution. Sci. Rep. 11, 6858. 10.1038/s41598-021-83664-5

Suetsugu, K., Ohta, T., Tayasu, I. (2018). Partial mycoheterotrophy in the leafless orchid *Cymbidium macrorhizon*. Am. J. Bot. 105(9): 1595–1600. 10.1002/ajb2.1142

Sun, Y., Moore, M.J., Lin, N., Adelalu, K.F., Meng, A., Jian, S., Yang, L., Li, J., Wang, H. (2017). Complete plastome sequencing of both living species of Circaeasteraceae (Ranunculales) reveals unusual rearrangements and the loss of the *ndh* gene family. BMC Genom. 9;18(1):592. 10.1186/s12864-017-3956-3

Tamura, K., Stecher, G., Kumar, S. (2021). MEGA11: Molecular volutionary Genetics Analysis Version 11 (FU Battistuzzi, Ed.). Mol. Biol. Evol. 38, 3022–3027. 10.1093/molbev/msab120

Thode, V.A., Lohmann, L.G. (2019). Comparative chloroplast genomics at low taxonomic levels: A case study using *Amphilophium* (Bignonieae, Bignoniaceae). Front. Plant Sci. 10, 796. 10.3389/fpls.2019.00796

Tu, X.D., Liu, D.K., Xu, S.W., Zhou, C.Y., Gao, X.Y., Zeng, M.Y., Zhang, S., Chen, J.L., Ma, L., Zhou, Z., Huang, M.Z., Chen, S.P., Liu, Z.J., Lan, S.R., Li, M.H. (2021). Plastid phylogenomics improves resolution of phylogenetic relationship in the *Cheirostylis* and *Goodyera* clades of Goodyerinae (Orchidoideae, Orchidaceae). Mol. Phyl. Evol. 164, 107269. 10.1016/j.ympev.2021.107269

Wen, Y., Qin, Y., Shao, B., Li, J., Ma, C., Liu, Y., Yang, B., Jin, X. (2022). The extremely reduced, diverged and reconfigured plastomes of the largest mycoheterotrophic orchid lineage. BMC Plant Biol. 22, 448 (2022). 10.1186/s12870-022-03836-x

WFO (2023). World Flora Online. Available at: http://www.worldfloraonline.org (Accessed in September 2023)

Wicke, S., Müller, K., De Pamphilis, C., Quandt, D. Bellot, S., Schneeweiss G (2016). Mechanistic model of evolutionary rate variation en route to a nonphotosynthetic lifestyle in plants. Proc. Nat. Acad. Sci. 113 (32), 9045–9050. 10.1073/pnas.1607576113

Wicke, S., Naumann, J. (2018). Molecular evolution of plastid genomes in parasitic flowering plants. Adv. Bot. Res. 85, 315–347. 10.1016/bs.abr.2017.11.014

Wu, S., Chen, J., Li, Y., Liu, A., Li, A., Yin, M., Shrestha, N., Liu, J., Ren, G. (2021). Extensive genomic rearrangements mediated by repetitive sequences in plastomes of *Medicago* and its relatives. BMC Plant Biol. 21, 421. 10.1186/s12870-021-03202-3

Yamori, W., Shikanai, T., Makino, A. (2015). Photosystem I cyclic electron flow via chloroplast NADH dehydrogenase-like complex performs a physiological role for photosynthesis at low light. Sci. Rep. 5, 13908. 10.1038/srep13908

Yang, J.B., Tang, M., Li, H.T., Zhang, Z.R., Li, D.Z. (2013). Complete chloroplast genome of the genus *Cymbidium*: lights into the species identification, phylogenetic implications and population genetic analyses. BMC Evol. Biol. 13, 84. 10.1186/1471-2148-13-84

Yule, G.U. (1925). A mathematical theory of evolution, based on the conclusions of Dr. J. C. Willis, F.R.S. Philos. Trans. R. Soc. B Biol. Sci. 213, 21–87. 10.1098/rstb.1925.0002

Zhang, G., Hu, Y., Huang, M.Z., Huang, W.C., Liu, D.K., Zhang, D., Hu, H., Downing, J.L., Liu, Z.J., Ma, H. (2023). Comprehensive phylogenetic analyses of Orchidaceae using nuclear genes and evolutionary insights into epiphytism. J. Integr. Plant Biol. 65(5), 1204–1225. 10.1111/jipb.13462

Zuckerkandl, E., Pauling, L. (1965). Molecules as documents of evolutionary history. J. Theor. Biol. 8, 357–366.

