## Supplementary Material 1 for "Plastid phylogenomics reveals evolutionary relationships in the mycoheterotrophic orchid genus *Dipodium* and provides insights into plastid gene degeneration"

**Table S1.1**: **Taxa included in phylogenetic analysis and divergence-time estimations.** Taxonomic concepts follow CHAH (2023) https://biodiversity.org.au/nsl/services/search/taxonomy for Australian taxa and WFO (2023): http://www.worldfloraonline.org for all other regions (accessed in May 2023) except for taxa with asterisks for which a synonym was used (shown in square brackets).

| Species | Subtribe | Tribe | Subfamily | NCBI/ ENA no. | DNA no. | Included in divergence-time analysis |
|---|---|---|---|---|---|---|
| *Aa paleacea* (Kunth) Rchb.f. | Cranichidiinae | Cranichideae | Orchidoideae | Givnish *et al.* (2015) | - | yes |
| *Acriopsis emarginata* D.L.Jones & M.A.Clem. | Cymbidiinae | Cymbidieae | Epidendroideae | This study | CNS_G00305 | yes |
| *Aganisia cyanea* (Lindl.) Rchb.f. | Zygopetalinae | Cymbidieae | Epidendroideae | Givnish *et al.* (2015) | - | yes |
| *Agapanthus coddii* F.M.Leight | - | - | Outgroup | NC 035971.1 | - | yes |
| *Agave americana* L. | - | - | Outgroup | NC 032053.1 | - | yes |
| *Angraecum sesquipedale* Thouars. | Angraecinae | Vandeae | Epidendroideae | Givnish *et al.* (2015) | - | yes |
| *Anoectochilus emeiensis* K.Y.Lang | Goodyerinae | Cranichideae | Orchidoideae | NC 033895.1 | - | yes |
| *Aphyllorchis montana* Rchb.f | - | Neottieae | Epidendroideae | NC 030703.1 | - | yes |
| *Apostasia odorata* Blume | - | - | Apostasioideae | NC 030722.1 | - | yes |
| *Apostasia shenzhenica* Z.J.Liu & L.J.Chen | - | - | Apostasioideae | NC 039812.1 | - | yes |
| *Apostasia wallichii* R.Br. | - | - | Apostasioideae | NC 036260.1 | - | yes |

| | | | | | | |
|---|---|---|---|---|---|---|
| *Arundina graminifolia* (D.Don) Hochr. | Arethusinae | Arethuseae | Epidendroideae | MN171408.1 | - | yes |
| *Asparagus officinalis* L. | - | - | Outgroup | NC 034777.1 | - | yes |
| *Batemannia colleyi* Lindl. | Zygopetalinae | Cymbidieae | Epidendroideae | Serna-Sánchez *et al.* (2021) | - | yes |
| *Bletilla striata* Rchb.f. | Coelogyninae | Arethuseae | Epidendroideae | NC 028422.1 | - | yes |
| *Brachionidium cruziae* L.O.Williams | Pleurothallidinae | Epidendreae | Epidendroideae | Serna-Sánchez *et al.* (2021) | - | yes |
| *Bulbophyllum inconspicuum* Maxim. | Dendrobiinae | Malaxideae | Epidendroideae | MN200377 | - | yes |
| *Calanthe aristulifera* Rchb.f. | - | Collabieae | Epidendroideae | MN200378 | - | yes |
| *Calanthe sylvatica* Lindl. | - | Collabieae | Epidendroideae | NC 044633.1 | - | yes |
| *Calopogon tuberosus* (L.) Britton, Sterns & Poggenb. | Arethusinae | Arethuseae | Epidendroideae | Givnish *et al.* (2015) | - | yes |
| *Calypso bulbosa* var. *occidentalis* (Holz.) Cockerell. | Calypsoinae | Epidendreae | Epidendroideae | NC 040980.1 | - | yes |
| *Catasetum integerrimum* Hook | Catasetinae | Cymbidieae | Epidendroideae | Givnish *et al.* (2015) | - | yes |
| *Cattleya liliputana* (Pabst) Van den Berg | Laeliinae | Epidendreae | Epidendroideae | NC 032083.1 | - | yes |
| *Cephalanthera damasonium* Druce | - | Neottieae | Epidendroideae | NC 041179.1 | - | yes |
| *Cephalanthera rubra* (L.) Rich. | - | Neottieae | Epidendroideae | NC 041181.1 | - | yes |
| *Changnienia amoena* S.S.Chien. | Calypsoinae | Epidendreae | Epidendroideae | NC 045402.1 | - | yes |
| *Chaubardia surinamensis* Rchb.f. | Zygopetalinae | Cymbidieae | Epidendroideae | Serna-Sánchez *et al.* (2021) | - | yes |
| *Cheirostylis chinensis* Rolfe | Goodyerinae | Cranichideae | Orchidoideae | MN641483.1 | - | yes |
| *Codonorchis lessonii* Lindl. | - | Codonorchideae | Orchidoideae | Givnish *et al.* (2015) | - | yes |
| *Coelogyne flaccida* Lindl. | Coelogyninae | Arethuseae | Epidendroideae | Givnish *et al.* (2015) | - | yes |
| *Corallorhiza striata* Lindl. | Coelogyninae | Arethuseae | Epidendroideae | NC 040978.1 | - | yes |
| *Corallorhiza trifida* Châtel | Coelogyninae | Arethuseae | Epidendroideae | NC 025662.1 | - | yes |
| *Coryanthes macrantha* Hook | Stanhopeinae | Cymbidieae | Epidendroideae | Givnish *et al.*(2015) | - | yes |
| *Cremastra appendiculata* (D.Don) Makino | Coelogyninae | Arethuseae | Epidendroideae | NC 037439.1 | - | yes |
| *Cremastra unguiculata* Finet. | Coelogyninae | Arethuseae | Epidendroideae | MN200381 | - | yes |
| *Cymbidium aloifolium* (L.) Sw. | Cymbidiinae | Cymbidieae | Epidendroideae | NC 021429.1 | - | yes |
| *Cymbidium canaliculatum* R.Br. | Cymbidiinae | Cymbidieae | Epidendroideae | This study | CNS_G00165 | yes |
| *Cymbidium ensifolium* Sw. | Cymbidiinae | Cymbidieae | Epidendroideae | NC 028525.1 | - | yes |
| *Cymbidium faberi* Rolfe | Cymbidiinae | Cymbidieae | Epidendroideae | NC 027743.1 | - | yes |
| *Cymbidium floribundum* Lindl. | Cymbidiinae | Cymbidieae | Epidendroideae | MN173778.1 | - | yes |
| *Cymbidium lancifolium* Hook | Cymbidiinae | Cymbidieae | Epidendroideae | NC 029712.1 | - | yes |

| | | | | | | |
|---|---|---|---|---|---|---|
| *Cymbidium macrorhizon* Lindl. | Cymbidiinae | Cymbidieae | Epidendroideae | NC 029713.1 | - | yes |
| *Cymbidium crassifolium* Wall. | Cymbidiinae | Cymbidieae | Epidendroideae | NC 021433.1 | - | yes |
| *Cypripedium calceolus* L. | - | - | Cypripedioideae | NC 045400.1 | - | yes |
| *Cypripedium japonicum* Thunb. | - | - | Cypripedioideae | NC 027227.1 | - | yes |
| *Cyrtopodium flavum* Link & Otto ex Rchb. | Cyrtopodiinae | Cymbidieae | Epidendroideae | Givnish *et al.* (2015) | - | yes |
| *Dendrobium kingianum* Bidwill ex Lindl. | Dendrobiinae | Malaxideae | Epidendroideae | LC331062.1 | - | yes |
| *Dendrobium moniliforme* (L.) Sw. | Dendrobiinae | Malaxideae | Epidendroideae | MN200384 | - | yes |
| *Dichaea pendula* (Aubl.) Cogn. | Zygopetalinae | Cymbidieae | Epidendroideae | Serna-Sánchez *et al.* (2021) | - | yes |
| *Dipodium* aff. *roseum** D.L.Jones & M.A.Clem. | Dipodiinae | Cymbidieae | Epidendroideae | This study | HTCG0828 | no |
| *Dipodium* aff. *roseum** D.L.Jones & M.A.Clem. | Dipodiinae | Cymbidieae | Epidendroideae | This study | HTCG0830 | no |
| *Dipodium* aff. *roseum** D.L.Jones & M.A.Clem. | Dipodiinae | Cymbidieae | Epidendroideae | This study | HTCG0831 | no |
| *Dipodium* aff. *roseum** D.L.Jones & M.A.Clem. | Dipodiinae | Cymbidieae | Epidendroideae | This study | HTCG0832 | yes |
| *Dipodium* aff. *stenocheilum** O.Schwarz | Dipodiinae | Cymbidieae | Epidendroideae | This study | HTCG1691 | no |
| *Dipodium ammolithum* M.D.Barrett, R.L.Barrett & K.W.Dixon | Dipodiinae | Cymbidieae | Epidendroideae | This study | HTCG1372 | yes |
| *Dipodium atropurpureum* D.L.Jones | Dipodiinae | Cymbidieae | Epidendroideae | This study | HTCG0760 | yes |
| *Dipodium atropurpureum* D.L.Jones | Dipodiinae | Cymbidieae | Epidendroideae | This study | HTCG1679 | no |
| *Dipodium basalticum* M.D.Barrett, R.L.Barrett & K.W.Dixon | Dipodiinae | Cymbidieae | Epidendroideae | This study | HTCG1693 | yes |
| *Dipodium campanulatum* D.L.Jones | Dipodiinae | Cymbidieae | Epidendroideae | This study | HTCG1680 | yes |
| *Dipodium campanulatum* D.L.Jones | Dipodiinae | Cymbidieae | Epidendroideae | This study | HTCG1681 | no |
| *Dipodium elegantulum* D.L.Jones | Dipodiinae | Cymbidieae | Epidendroideae | This study | HTCG1682 | yes |
| *Dipodium ensifolium* F.Muell. | Dipodiinae | Cymbidieae | Epidendroideae | This study | HTCG1343 | yes |
| *Dipodium hamiltonianum* F.M.Bailey | Dipodiinae | Cymbidieae | Epidendroideae | This study | HTCG1683 | yes |
| *Dipodium hamiltonianum** F.M.Bailey [syn. *Dipodium interaneum* D.L.Jones] | Dipodiinae | Cymbidieae | Epidendroideae | This study | HTCG0181 | yes |
| *Dipodium pandanum* F.M.Bailey | Dipodiinae | Cymbidieae | Epidendroideae | This study | CNS_G01262 | yes |
| *Dipodium pandanum* F.M.Bailey | Dipodiinae | Cymbidieae | Epidendroideae | This study | HTCG1694 | no |
| *Dipodium pardalinum* D.L.Jones | Dipodiinae | Cymbidieae | Epidendroideae | This study | HTCG1684 | yes |
| *Dipodium pardalinum* D.L.Jones | Dipodiinae | Cymbidieae | Epidendroideae | This study | HTCG1685 | no |
| *Dipodium punctatum** (Sm.) R.Br. [syn. *Dipodium pulchellum* D.L.Jones & M.A.Clem.] | Dipodiinae | Cymbidieae | Epidendroideae | This study | HTCG1686 | yes |
| *Dipodium punctatum* (Sm.) R.Br. | Dipodiinae | Cymbidieae | Epidendroideae | This study | HTCG0827 | yes |
| *Dipodium roseum* D.L.Jones & M.A.Clem. | Dipodiinae | Cymbidieae | Epidendroideae | MN200386 | - | no |

| | | | | | | |
|---|---|---|---|---|---|---|
| *Dipodium roseum* D.L.Jones & M.A.Clem. | Dipodiinae | Cymbidieae | Epidendroideae | This study | HTCG1687 | yes |
| *Dipodium roseum* D.L.Jones & M.A.Clem. | Dipodiinae | Cymbidieae | Epidendroideae | This study | HTCG1688 | no |
| *Dipodium stenocheilum* O.Schwarz | Dipodiinae | Cymbidieae | Epidendroideae | This study | HTCG1689 | no |
| *Dipodium stenocheilum* O.Schwarz | Dipodiinae | Cymbidieae | Epidendroideae | This study | HTCG1690 | yes |
| *Dipodium variegatum* M.A.Clem. & D.L.Jones | Dipodiinae | Cymbidieae | Epidendroideae | This study | HTCG1692 | yes |
| *Elleanthus sodiroi* Schltr. | - | Sobralieae | Epidendroideae | NC 027266.1 | - | yes |
| *Epipactis mairei* Schltr. | - | Neottieae | Epidendroideae | NC 030705.1 | - | yes |
| *Epipactis thunbergia* A.Gray. | - | Neottieae | Epidendroideae | MN200387 | - | yes |
| *Epipactis veratrifolia* Boiss. & [illegible] | - | Neottieae | Epidendroideae | NC 030708.1 | - | yes |
| *Eria scabrilinguis* Lindl. | - | Podochileae | Epidendroideae | MN477202.1 | - | yes |
| *Eulophia bicallosa* (D.Don) P.F.Hunt & Summerh. | Eulophiinae | Cymbidieae | Epidendroideae | This study | HTCG1696 | yes |
| *Eulophia graminea* Lindl. | Eulophiinae | Cymbidieae | Epidendroideae | This study | CNS_G02766 | yes |
| *Eulophia nuda* Lindl. | Eulophiinae | Cymbidieae | Epidendroideae | This study | HTCG1697 | yes |
| *Eulophia petersii* Rchb.f. | Eulophiinae | Cymbidieae | Epidendroideae | Givnish *et al.* (2015) | - | no |
| *Evotella carnosa* (Lindl.) J.C.Manning & Goldblatt | Coryciinae | Orchideae | Orchidoideae | Givnish *et al.* (2015) | - | yes |
| *Gastrochilus japonicus* Schltr. | Aeridinae | Vandeae | Epidendroideae | NC 035833.1 | - | yes |
| *Geodorum densiflorum* (Lam.) Schltr. | Eulophiinae | Cymbidieae | Epidendroideae | This study | CNS_G01890 | yes |
| *Gongora pleiochroma* Rchb.f. | Stanhopeinae | Cymbidieae | Epidendroideae | Givnish *et al.* (2015) | - | yes |
| *Goodyera fumata* Thwaites. | Goodyerinae | Cranichideae | Orchidoideae | NC 026773.1 | - | yes |
| *Goodyera schlechtendaliana* Rchb.f. | Goodyerinae | Cranichideae | Orchidoideae | MK134679.1 | - | yes |
| *Guarianthe aurantiaca* (Bateman) Dressler & W.E.Higgins. | Laeliinae | Epidendreae | Epidendroideae | Givnish *et al.* (2015) | - | yes |
| *Gymnadenia conopsea* (L.) R.Br. | Orchidinae | Orchideae | Orchidoideae | MN200391 | - | yes |
| *Habenaria arenaria* Lindl. | Orchidinae | Orchideae | Orchidoideae | Givnish *et al.* (2015) | - | yes |
| *Habenaria ciliolaris* Kraenzl. | Orchidinae | Orchideae | Orchidoideae | MN495954.1 | - | yes |
| *Hexalectris warnockii* Ames & Correll. | Laeliinae | Epidendreae | Epidendroideae | MH444822.1 | - | yes |
| *Holcoglossum lingulatum* (Aver.) Aver. | Aeridinae | Vandeae | Epidendroideae | NC 041465.1 | - | yes |
| *Holcoglossum subulifolium* (Rchb.f.) Christenson. | Aeridinae | Vandeae | Epidendroideae | NC 041519.1 | - | yes |
| *Huntleya meleagris* Lindl. | Zygopetalinae | Cymbidieae | Epidendroideae | Serna-Sánchez *et al.* (2021) | - | yes |
| *Iris sanguinea* Hornem | - | - | Outgroup | NC 029227.1 | - | yes |
| *Lilium pensylvanicum* Ker Gawl. | - | - | Outgroup | NC 043876.1 | - | yes |

| | | | | | | |
|---|---|---|---|---|---|---|
| *Liparis auriculata* Blume ex Miq. | Malaxidinae | Malaxideae | Epidendroideae | MN200365 | - | yes |
| *Ludisia discolor* (Ker Gawl.) Blume | Goodyerinae | Cranichideae | Orchidoideae | NC 030540.1 | - | yes |
| *Masdevallia coccinea* Linden ex Lindl. | Pleurothallidinae | Epidendreae | Epidendroideae | NC 026541.1 | - | yes |
| *Maxillaria nasuta* Rchb.f. | Maxillariinae | Cymbidieae | Epidendroideae | Givnish *et al.* (2015) | - | yes |
| *Maxillaria sanderiana* Rchb.f. ex Sander | Maxillariinae | Cymbidieae | Epidendroideae | Givnish *et al.* (2015) | - | yes |
| *Neottia listeroides* Lindl. | - | Neottieae | Epidendroideae | NC_030713.1 | - | no |
| *Neottia ovata* Bluff & Fingerh. | - | Neottieae | Epidendroideae | NC 030712.1 | - | yes |
| *Nervilia simplex* (Spreng.) Schltr. | - | Nervilieae | Epidendroideae | Givnish *et al.* (2015) | - | no |
| *Neuwiedia zollingeri* var. *singapureana* (Wall. ex Baker) de Vogel | - | - | Apostasioideae | LC199503.1 | - | yes |
| *Oberonia japonica* (Maxim.) Makino | Malaxidinae | Malaxideae | Epidendroideae | NC 035832.1 | - | yes |
| *Oeceoclades pelorica* (D.L.Jones & M.A.Clem.) D.L.Jones & M.A.Clem. | Eulophiinae | Cymbidieae | Epidendroideae | This study | HTCG1695 | yes |
| *Oncidium sphacelatum* Lindl. | Oncidiinae | Cymbidieae | Epidendroideae | NC 028148.1 | - | yes |
| *Ophrys fusca* subsp. *iricolor** (Desf.) K.Richt. | Orchidinae | Orchideae | Orchidoideae | AP018716.1 | - | yes |
| *Ophrys sphegodes* Mill. | Orchidinae | Orchideae | Orchidoideae | AP018717.1 | - | yes |
| *Otoglossum globuliferum* (Kunth) N.H.Williams & M.W.Chase | Zygopetalinae | Cymbidieae | Epidendroideae | Serna-Sánchez *et al.* (2021) | - | yes |
| *Otostylis brachystali* Schltr. | Zygopetalinae | Cymbidieae | Epidendroideae | Serna-Sánchez *et al.* (2021) | - | yes |
| *Pabstia jugosa* (Lindl.) Garay. | Zygopetalinae | Cymbidieae | Epidendroideae | Serna-Sánchez *et al.* (2021) | - | yes |
| *Palmorchis pabstii* Veyret. | - | Neottieae | Epidendroideae | NC 041190.1 | - | yes |
| *Paphiopedilum armeniacum* S.C.Chen & F.Y.Liu. | - | - | Cypripedioideae | NC 026779.1 | - | yes |
| *Paphiopedilum niveum* (Rchb.f.) Stein | - | - | Cypripedioideae | NC 026776.1 | - | yes |
| *Pelatantheria scolopendrifolia* (Makino) Aver. | Aeridinae | Vandeae | Epidendroideae | NC 035829.1 | - | yes |
| *Pescatoria wallisii* Linden & Rchb.f. | Zygopetalinae | Cymbidieae | Epidendroideae | Serna-Sánchez *et al.* (2021) | - | yes |
| *Phaius tankervilleae* (Banks) Blume. | - | Collabieae | Epidendroideae | Givnish *et al.* (2015) | - | yes |
| *Phalaenopsis japonica* (Rchb.f.) Kocyan & Schuit. | Aeridinae | Vandeae | Epidendroideae | Givnish *et al.* (2015) | - | yes |
| *Phalaenopsis pulcherrima* (Lindl.) J.J.Sm. | Aeridinae | Vandeae | Epidendroideae | MG459020.1 | - | yes |
| *Phragmipedium longifolium* (Rchb.f. & Warsz.) Rolfe | - | - | Cypripedioideae | NC 028149.1 | - | yes |
| *Platanthera mandarinorum* Rchb.f. | Orchidinae | Orchideae | Orchidoideae | MN200370 | - | yes |
| *Platystele aurea* Garay. | Pleurothallidinae | Epidendreae | Epidendroideae | Serna-Sánchez *et al.* (2021) | - | yes |
| *Pleione formosana* Hayata | Coelogyninae | Arethuseae | Epidendroideae | NC 042197.1 | - | yes |
| *Pogonia ophioglossoides* (L.) Ker Gawl. | - | Pogonieae | Vanilloideae | Givnish *et al.* (2015) | - | yes |

| | | | | | | |
|---|---|---|---|---|---|---|
| *Ponerorchis gracilis* (Blume) X.H.Jin, Schuit. & W.T.Jin | Orchidinae | Orchideae | Orchidoideae | MN200376 | - | yes |
| *Scaphosepalum antenniferum* Rolfe. | Pleurothallidinae | Epidendreae | Epidendroideae | Serna-Sánchez *et al.* (2021) | - | yes |
| *Sobralia mucronate* Ames & C.Schweinf. | - | Sobralieae | Epidendroideae | Givnish *et al.* (2015) | - | yes |
| *Spiranthes sinensis* (Pers.) Ames | Spiranthinae | Cranichideae | Orchidoideae | MK936427.1 | - | yes |
| *Teagueia* (Luer) Luer. | Pleurothallidinae | Epidendreae | Epidendroideae | Serna-Sánchez *et al.* (2021) | - | yes |
| *Telipogon glicensteinii* Dodson & R.Escobar | Oncidiinae | Cymbidieae | Epidendroideae | Givnish *et al.* (2015) | - | yes |
| *Thelymitra cyanea* (Lindl.) Benth. | Thelymitrinae | Diurideae | Orchidoideae | Givnish *et al.* (2015) | - | yes |
| *Thrixspermum japonicum* Rchb.f. | Aeridinae | Vandeae | Epidendroideae | NC 035831.2 | - | yes |
| *Triphora trianthophora* (Sw.) Rydb. | Triphorinae | Triphoreae | Epidendroideae | Givnish *et al.* (2015) | - | no |
| *Tropidia polystachya* Ames. | - | Tropidieae | Epidendroideae | Givnish *et al.* (2015) | - | yes |
| *Vanda brunnea* Rchb.f. | Aeridinae | Vandeae | Epidendroideae | NC 041522.1 | - | yes |
| *Vanda falcata* Beer. | Aeridinae | Vandeae | Epidendroideae | NC 036372.1 | - | |
| *Vanilla aphylla* Blume | - | Vanilleae | Vanilloideae | NC 035320.1 | - | yes |
| *Vanilla planifolia* Andrews | - | Vanilleae | Vanilloideae | NC 026778.1 | - | yes |
| *Vanilla pompona* Schiede | - | Vanilleae | Vanilloideae | NC 036809.1 | - | yes |
| *Zootrophion hirtzii* Luer. | Pleurothallidinae | Epidendreae | Epidendroideae | Serna-Sánchez *et al.* (2021) | - | yes |
| *Zygopetalum triste* Barb.Rodr. | Zygopetalinae | Cymbidieae | Epidendroideae | Serna-Sánchez *et al.* (2021) | - | yes |
