## Supplementary Material 3 for "Plastid phylogenomics reveals evolutionary relationships in the mycoheterotrophic orchid genus *Dipodium* and provides insights into plastid gene degeneration"

ML-Phylogenetic tree of Orchidaceae.

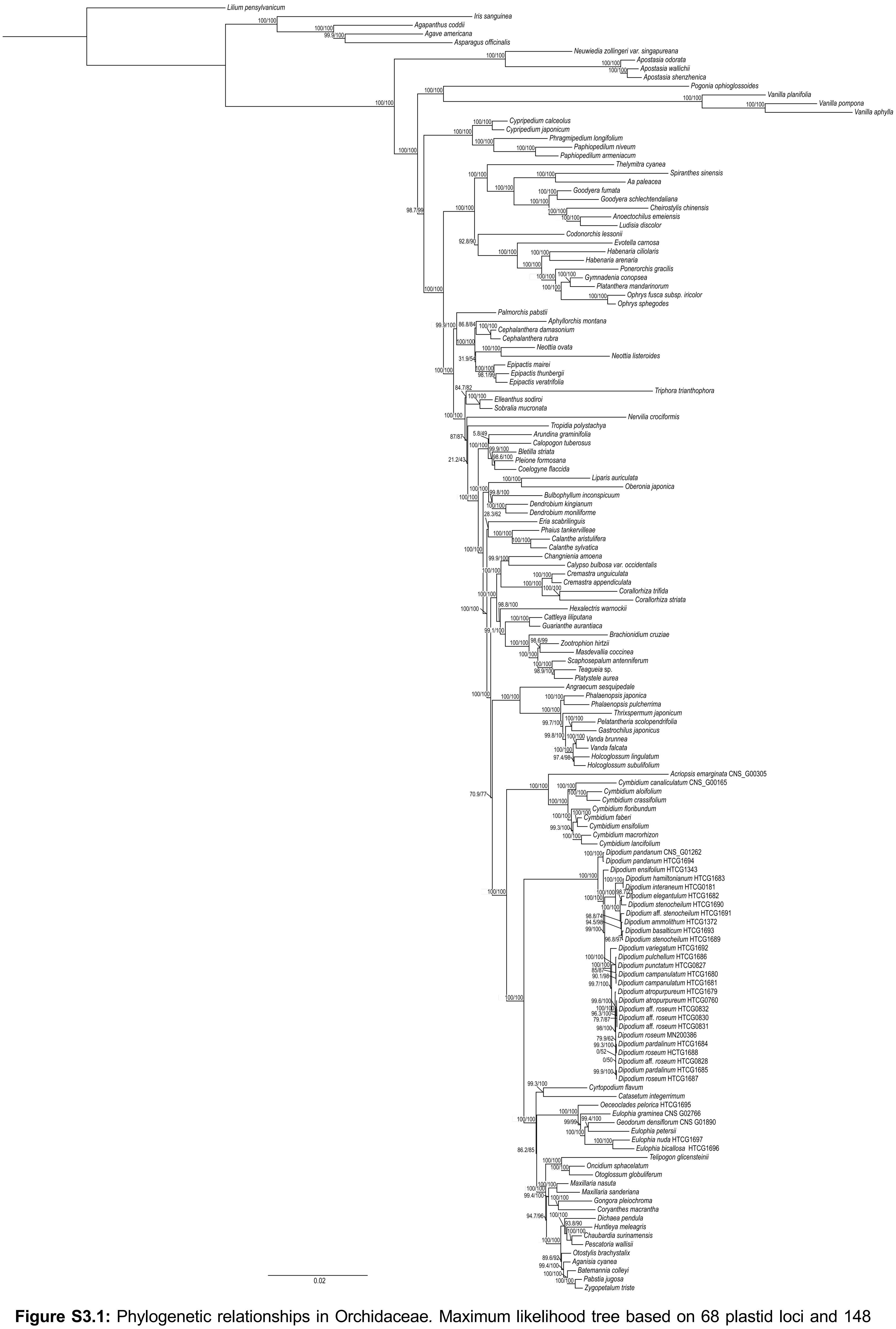


**Figure S3.1:** Phylogenetic relationships in Orchidaceae. Maximum likelihood tree based on 68 plastid loci and 148 taxa. Support values are given above each branch, SHaLRT is followed by UFBoot values. Scale bar represents branch length, along which 0.02 per-site substitutions are expected.
