## Supplementary Material 4 for "Plastid phylogenomics reveals evolutionary relationships in the mycoheterotrophic orchid genus *Dipodium* and provides insights into plastid gene degeneration"

**Table S4.1: Model Comparison by AICM** (Akaike Information Criterion by MCMC), evaluated with the model selection criterion by AIC of Fabozzi et al., (2014). Best fit model for the data set after AICM is the optimised relaxed clock, birth-death model (in bold).

| Model | AICM likelihood | +/- | burn-in | bootstrap replicates | delta AICM |
|---|---|---|---|---|---|
| **Optimised relaxed clock/Birth-death** | **432312.4507** | **2.5475** | **10** | **1000** | **0** |
| Optimised relaxed clock/ Yule | 432319.0708 | 2.5574 | 10 | 1000 | 6.6201 |
| Strict clock/ Birth-death | 437771.8470 | 1.6389 | 10 | 1000 | 5459.3963 |
| Strict clock/ Yule | 437764.7433 | 1.5339 | 10 | 1000 | 5452.2926 |

**From an article (appendix) titled:** ***Model Selection Criterion: AIC and BIC***:
*The Basics of Financial Econometrics: Tools, Concepts, and Asset Management Applications.* Frank J. Fabozzi, Sergio M. Focardi, Svetlozar T. Rachev and Bala G. Arshanapalli. © 2014 John Wiley & Sons, Inc. Published 2014 by John Wiley & Sons, Inc.

**Table S4.2: Comparison divergence-time estimations of major Orchidaceae linages (subfamilies), the tribe Cymbidieae and subtribe Dipodiinae**. Best fit model for the data set after AICM is the optimised relaxed clock, birth-death model (in bold). Absolute ages in 'million years ago' (Ma). MRCA: Most common recent ancestor; HDP: Highest posterior density interval.

| | **relaxed clock, Birth-death** | | | relaxed clock, Yule | | |
|---|---|---|---|---|---|---|
| MRCA | **mean ages** | **95% HDP lower bound** | **95% HDP upper bound** | mean ages | 95% HDP lower bound | 95% HDP upper bound |
| Orchidaceae | **126.35** | **100.56** | **155.60** | 124.24 | 100.49 | 152.42 |
| Apostasioideae | **101.96** | **98.01** | **105.82** | 101.96 | 98.18 | 105.95 |
| Vanilloideae | **93.26** | **89.27** | **97.05** | 93.23 | 89.38 | 97.13 |
| Cypripedioideae | **88.33** | **84.44** | **92.37** | 88.30 | 84.33 | 92.37 |
| Orchidoideae | **77.71** | **74.15** | **81.52** | 77.71 | 73.83 | 81.22 |
| Epidendroideae | **77.71** | **74.15** | **81.52** | 77.71 | 73.83 | 81.22 |
| Cymbidieae | **42.24** | **34.25** | **50.11** | 42.44 | 34.81 | 50.18 |
| Dipodiinae | **33.33** | **26.41** | **40.59** | 33.60 | 26.46 | 40.47 |

| | strict clock, Birth-death | | | strict clock, Yule | | |
|---|---|---|---|---|---|---|
| MRCA | mean ages | 95% HDP lower bound | 95% HDP upper bound | mean ages | 95% HDP lower bound | 95% HDP upper bound |
| Orchidaceae | 140.06 | 134.79 | 145.77 | 139.97 | 134.67 | 145.65 |
| Apostasioideae | 106.69 | 103.73 | 109.70 | 106.65 | 103.62 | 109.65 |
| Vanilloideae | 100.26 | 97.24 | 103.26 | 100.21 | 97.17 | 103.15 |
| Cypripedioideae | 80.44 | 77.69 | 83.24 | 80.48 | 77.69 | 83.28 |
| Orchidoideae | 71.20 | 68.86 | 73.65 | 71.24 | 68.84 | 73.62 |
| Epidendroideae | 71.20 | 68.86 | 73.65 | 71.24 | 68.84 | 73.62 |
| Cymbidieae | 44.80 | 42.52 | 46.52 | 44.92 | 42.83 | 46.48 |
| Dipodiinae | 36.51 | 34.81 | 38.20 | 36.57 | 34.92 | 38.30 |
