## Supplementary Material 5 for "Plastid phylogenomics reveals evolutionary relationships in the mycoheterotrophic orchid genus *Dipodium* and provides insights into plastid gene degeneration"

Maximum-clade-credibility tree from Bayesian divergence-time estimations of Orchidaceae.

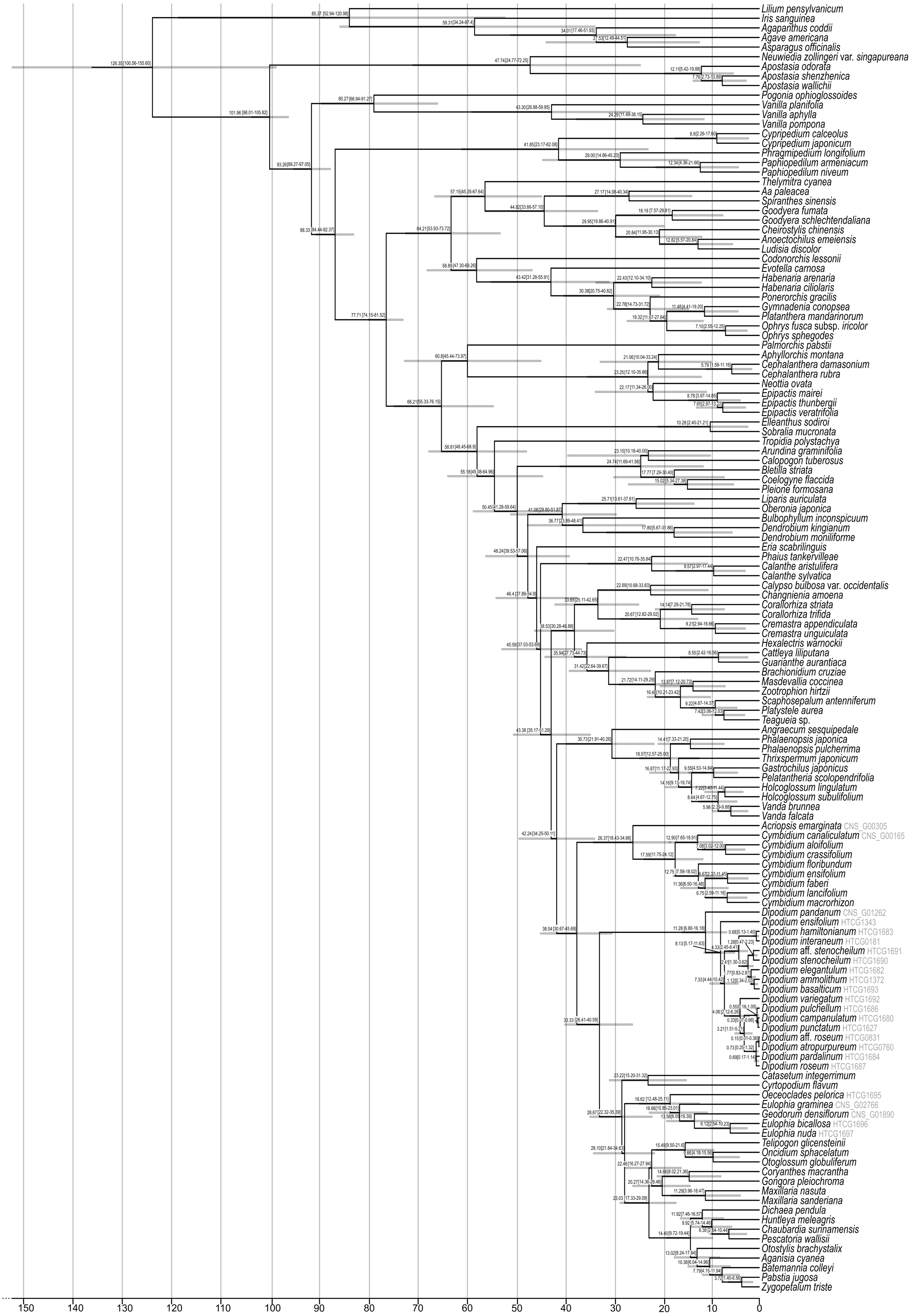


**Figure S5.1: Chronogram of Orchidaceae.** Maximum-clade-credibility tree from Bayesian divergence-time estimation in BEAST2 based on 30 plastid loci, 134T and an optimised relaxed molecular clock model under the birth-death prior. Divergence times (million years ago) are shown at each node, together with 95% highest posterior density (HDP) values indicated by grey bars and values in parentheses.
