## Supplementary Material 6 for "Plastid phylogenomics reveals evolutionary relationships in the mycoheterotrophic orchid genus *Dipodium* and provides insights into plastid gene degeneration"

**Table S6.1**: Summery of the assembly features of 24 *Dipodium* plastomes reconstructed in this study.

| | Samlpe | # trimmed reads | # mapped reads | mean coverage |
|---|---|---|---|---|
| 1 | *Dipodium ammolithum* HTCG1372 | 8,131,829 | 51,851 | 63 |
| 2 | *Dipodium atropurpureum* HTCG0760 | 3,802,046 | 114,178 | 166 |
| 3 | *Dipodium atropurpureum* HTCG1679 | 5,293,802 | 76,117 | 71 |
| 4 | *Dipodium basalticum* HTCG1693 | 7,476,898 | 56,106 | 73 |
| 5 | *Dipodium campanulatum* HTCG1680 | 4,928,715 | 33,297 | 31 |
| 6 | *Dipodium campanulatum* HTCG1681 | 4,714,583 | 65,257 | 75 |
| 7 | *Dipodium elegantulum* HTCG1682 | 5,696,744 | 40,461 | 48 |
| 8 | *Dipodium ensifolium* HTCG1343 | 11,895,622 | 483,386 | 627 |
| 9 | *Dipodium hamiltonianum* HTCG1683 | 11,327,386 | 54,724 | 63 |
| 10 | *Dipodium interaneum* HTCG0181 | 3,709,623 | 53,624 | 75 |
| 11 | *Dipodium pandanum* CNS_G01262 | 332,604 | 89,782 | 204 |
| 12 | *Dipodium pardalinum* HTCG1684 | 8,086,850 | 70,697 | 67 |
| 13 | *Dipodium pardalinum* HTCG1685 | 27,999,734 | 201,257 | 198 |
| 14 | *Dipodium pulchellum* HTCG1686 | 6,795,229 | 85,282 | 93 |
| 15 | *Dipodium punctatum* HTCG0827 | 8,987,460 | 90,899 | 120 |
| 16 | *Dipodium roseum* HTCG1687 | 7,399,306 | 90,802 | 101 |
| 17 | *Dipodium roseum* HTCG1688 | 6,207,464 | 52,233 | 50 |
| 18 | *Dipodium* aff. *roseum* HTCG0828 | 7,953,368 | 251,830 | 353 |
| 19 | *Dipodium* aff. *roseum* HTCG0830 | 4,347,368 | 56,127 | 74 |
| 20 | *Dipodium* aff. *roseum* HTCG0831 | 11,388,084 | 357,584 | 475 |
| 21 | *Dipodium* aff. *roseum* HTCG0832 | 10,706,617 | 244,503 | 336 |
| 22 | *Dipodium stenocheilum* HTCG1689 | 4,657,080 | 38,117 | 44 |
| 23 | *Dipodium stenocheilum* HTCG1690 | 6,026,488 | 37,602 | 47 |
| 24 | *Dipodium variegatum* HTCG1692 | 4,256,510 | 41,813 | 44 |
