## Supplementary Material 7 for "Plastid phylogenomics reveals evolutionary relationships in the mycoheterotrophic orchid genus *Dipodium* and provides insights into plastid gene degeneration"

Circular plastome maps of 24 newly generated *Dipodium* plastomes.

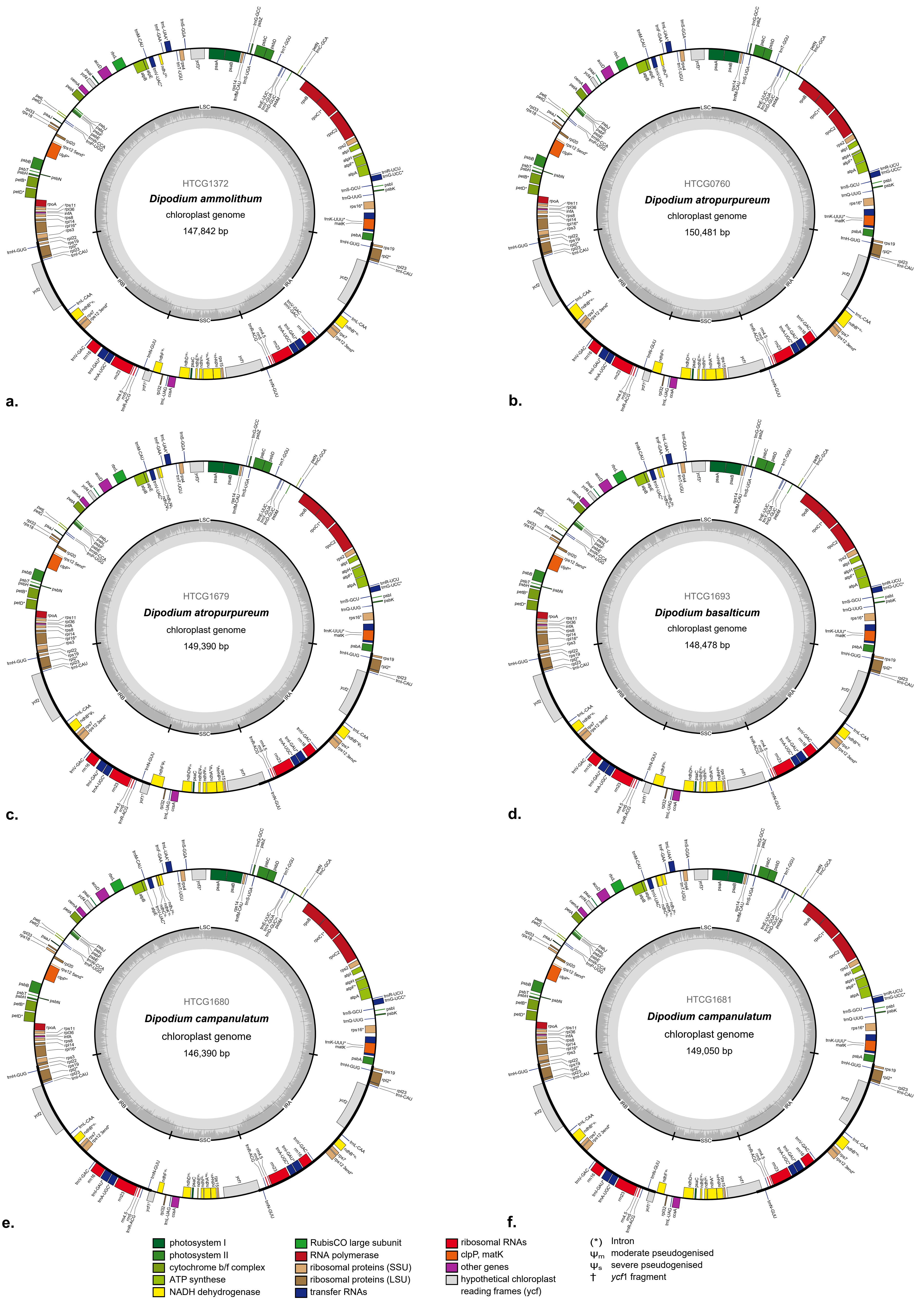


**Figure S7.1**: Circular plastome maps of a. *D. ammolithum* (HTCG1372), b., c. *D. atropurpureum* (HTCG0760, HTCG1679), d. *D. basalticum* (HTCG1683) and e., f. *D. campanulatum* (HTCG1680, HTCG1681). Genes outside the circle are transcribed in a clockwise direction, those inside the circle are transcribed in a counterclockwise direction. The dark grey inner circle corresponds to the G/C content, and the lighter grey to the A/C content. SSC: Small Single Copy; LSC: Large Single Copy: IRA/B: Inverted Repeat A/B.

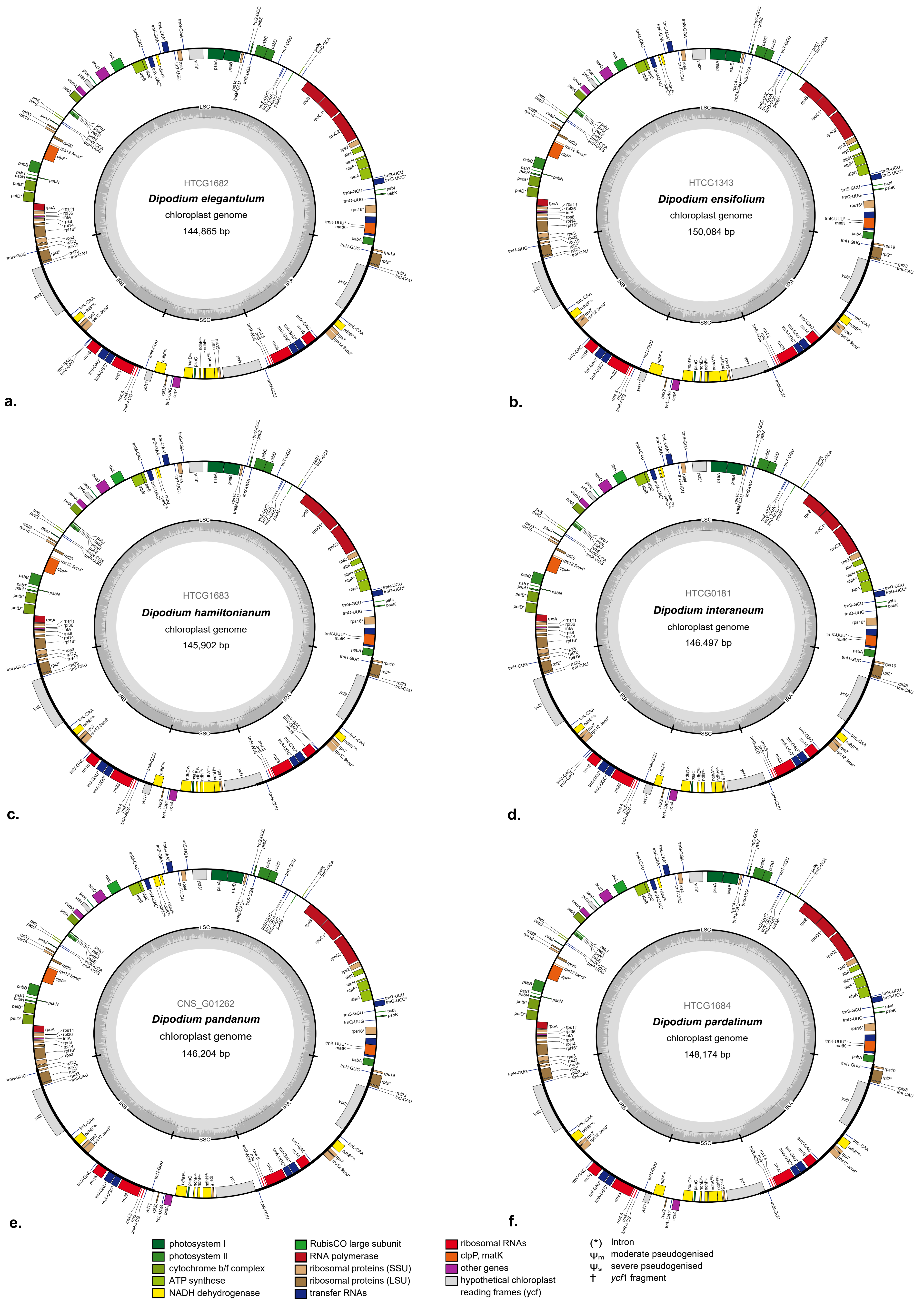


**Figure S7.2**: Circular plastome maps of a. *D. elegantulum* (HTCG1682), b. *D. ensifolium* (HTCG1343), c. *D. hamiltonianum* (HTCG1683), d. *D. interaneum* (HTCG0181), e. *D. pandanum* (CNS_G01262) and f. *D. pardalinum* (HTCG1684). Genes outside the circle are transcribed in a clockwise direction, those inside the circle are transcribed in a counterclockwise direction. The dark grey inner circle corresponds to the G/C content, and the lighter grey to the A/C content. SSC: Small Single Copy; LSC: Large Single Copy: IRA/B: Inverted Repeat A/B.

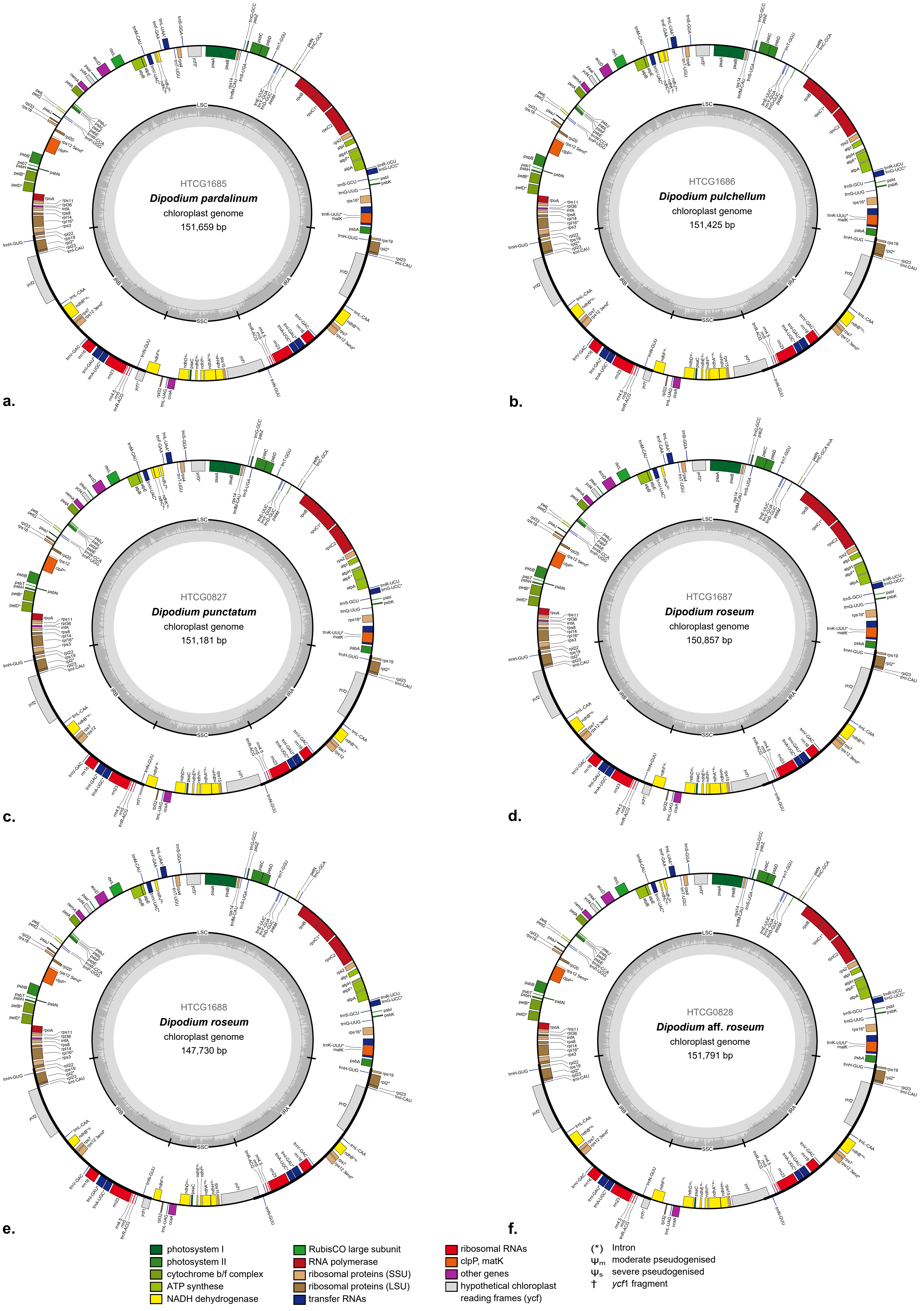


**Figure S7.3**: Circular plastome maps of a. *D. pardalinum* (HTCG1685), b. *D. pulchellum* (HTCG1686), c. *D. punctatum* (HTCG0827), d., e. *D. roseum* (HTCG1687, HTCG1688) and f. *D.* aff. *roseum* (HTCG0828). Genes outside the circle are transcribed in a clockwise direction, those inside the circle are transcribed in a counterclockwise direction. The dark grey inner circle corresponds to the G/C content, and the lighter grey to the A/C content. SSC: Small Single Copy; LSC: Large Single Copy: IRA/B: Inverted Repeat A/B.

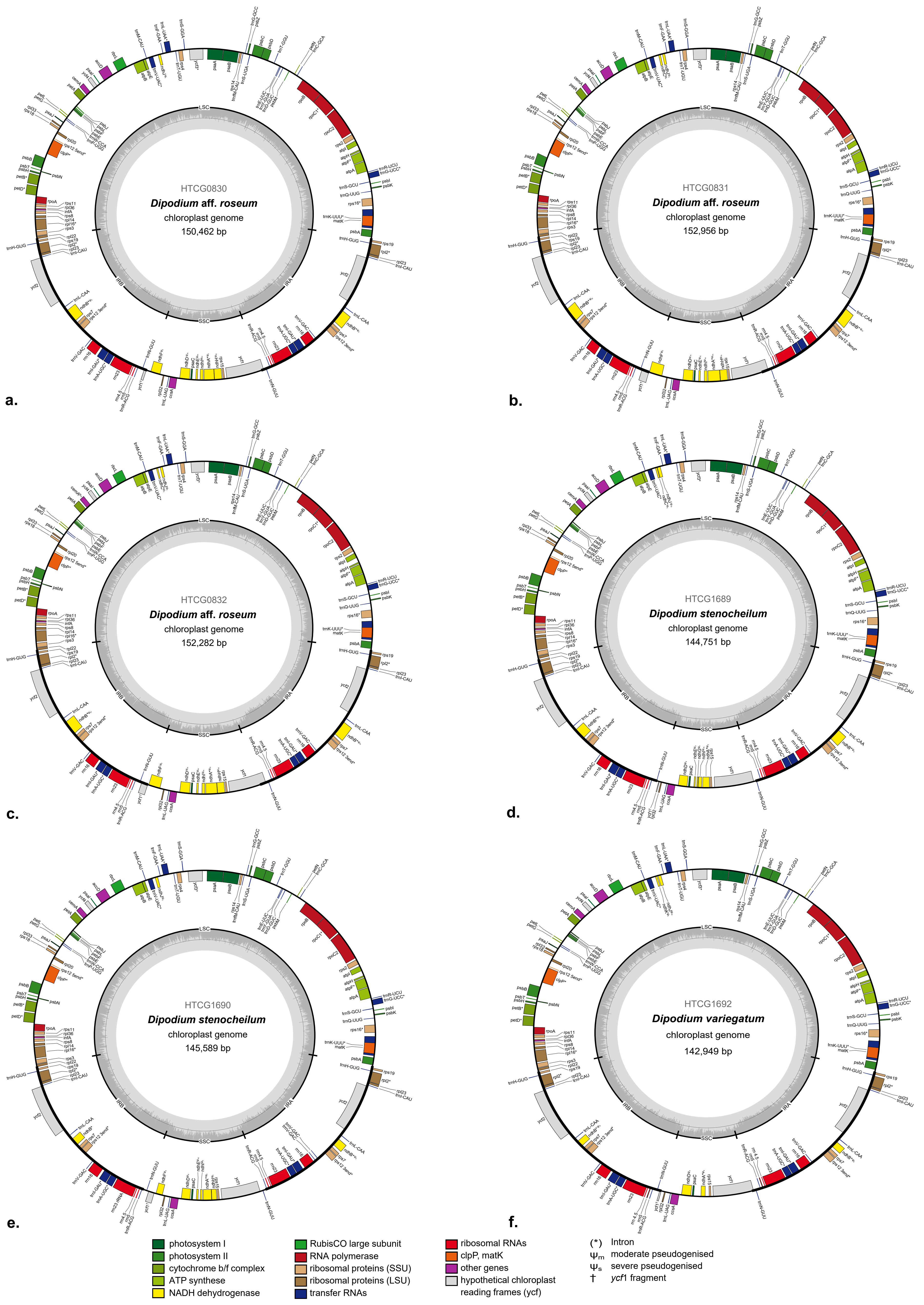


**Figure S7.4**: Circular plastome maps of a.-c. *D.* aff. *roseum* (HTCG0830, HTCG0831, HTCG0832), d., e. *D. stenocheilum* (HTCG1689, HTCG1690) and f. *D. variegatum* (HTCG1692) .Genes outside the circle are transcribed in a clockwise direction, those inside the circle are transcribed in a counterclockwise direction. The dark grey inner circle corresponds to the G/C content, and the lighter grey to the A/C content. SSC: Small Single Copy; LSC: Large Single Copy: IRA/ B: Inverted Repeat A/B.
